# “Off-pore” nucleoporins relocalize heterochromatic breaks through phase separation

**DOI:** 10.1101/2023.12.07.570729

**Authors:** Chiara Merigliano, Taehyun Ryu, Colby D. See, Christopher P. Caridi, Xiao Li, Nadejda L. Butova, Trevor W. Reynolds, Changfeng Deng, David M. Chenoweth, Maya Capelson, Irene Chiolo

## Abstract

Phase separation forms membraneless compartments in the nuclei, including by establishing heterochromatin “domains” and repair foci. Pericentromeric heterochromatin mostly comprises repeated sequences prone to aberrant recombination, and “safe” homologous recombination (HR) repair of these sequences requires the movement of repair sites to the nuclear periphery before Rad51 recruitment and strand invasion. How this mobilization initiates is unknown, and the contribution of phase separation to these dynamics is unclear. Here, we show that Nup98 nucleoporin is recruited to heterochromatic repair sites before relocalization through Sec13 or Nup88 nucleoporins, and downstream from the Smc5/6 complex and SUMOylation. Remarkably, the phase separation properties of Nup98 are required and sufficient to mobilize repair sites and exclude Rad51, thus preventing aberrant recombination while promoting HR repair. Disrupting this pathway results in heterochromatin repair defects and widespread chromosome rearrangements, revealing a novel “off-pore” role for nucleoporins and phase separation in nuclear dynamics and genome integrity in a multicellular eukaryote.

**Highlights:**

- Nup88 and Sec13 recruit Nup98 to heterochromatic DSBs downstream from Smc5/6
- Nup88, Sec13 and Nup98 promote repair focus mobilization in heterochromatin ‘off-pore’
- Nup98 excludes Rad51 from repair sites inside the heterochromatin domain
- Phase separation by Nup98 is required and sufficient for relocalization and Rad51 exclusion

## INTRODUCTION

Heterochromatin comprises ∼10-30% of human and fly genomes^1^, is mostly composed of repeated DNA sequences (*e.g.,* transposons, and ‘satellite’ repeats^2,3^), and is typically characterized by ‘silent’ epigenetic marks (*e.g.*, H3K9me2/3 and HP1^4^). In heterochromatin, thousands to millions of identical repeated sequences, including those from different chromosomes, can engage in ectopic recombination, presenting a serious threat to genome stability in multicellular eukaryotes^5^. Specialized mechanisms promote homologous recombination (HR) repair in pericentromeric heterochromatin (hereafter ‘heterochromatin’) while preventing aberrant recombination^6–9^.

In *Drosophila* cells, where heterochromatin forms a distinct nuclear domain^4,10,11^, “safe” HR repair relies on the relocalization of double-strand breaks (DSBs) to the nuclear periphery before Rad51 recruitment and strand invasion^10,12,13^. Relocalization likely isolates damaged DNA from similar repeated sequences on non-homologous chromosomes, thus promoting ‘safe’ exchanges with sister chromatids or homologs^6,7,10,12–16^. A similar relocalization occurs in mouse cells^13,17–19^, demonstrating conserved pathways. Relocalization and the initial block to HR progression require SUMOylation by the SUMO E3-ligases dPIAS and the Smc5/6 complex^10,12,13,15^. At the nuclear periphery, the nucleoporin Nup107 mediates anchoring of repair sites to the nuclear pores, and this association is required for restarting repair *via* Brca2 and Rad51 recruitment^12^.

Key components of this pathway also include a transient network of nuclear actin filaments (F-actin) formed by Arp2/3 at repair sites, which mediate nuclear myosin-driven relocalization through directed motions^13^. Notably, filaments and directed motions are mostly detected between the heterochromatin domain periphery and the nuclear periphery^13^, raising the question of how heterochromatic breaks reach the heterochromatin domain periphery in the first place.

While the movement of heterochromatic repair sites occur in the context of phase separated structures, including repair foci^20,21^, the heterochromatin domain^22,23^, and nucleoporins^24–27^, how phase separation contributes to these dynamics is unknown. Nuclear pore proteins also found in the nucleoplasm, where they regulate transcription during *Drosophila* development^28–31^ and promote leukemic transformation in chimeric fusions resulting from translocations^32–34^. Whether these components contribute to repair “off pore” is unclear, and a role for nucleoporin-mediated phase separation in this context has not been investigated. Here, we establish that the nucleoporin Nup98, a prime determinant of leukemic transformation, is recruited to heterochromatic repair foci in the nucleoplasm downstream from SUMOylation, and its phase separation properties are required and sufficient for the mobilization of heterochromatic repair sites and Rad51 exclusion.

## RESULTS

### Nup88, Nup98 and Sec13 are required for relocalizing heterochromatic DSBs

In *Drosophila* cells, repair sites leave the heterochromatin domain starting 10 min after DSB induction with ionizing radiation (IR), resulting in fewer repair sites (γH2Av foci) in DAPI-bright heterochromatin 60 min after IR^10,12,13^. Defective anchoring to the nuclear periphery after Nup107 RNAi depletion results in break sites returning to the heterochromatin domain, leading to a higher number of γH2Av foci in DAPI-bright 60 min after IR without affecting the total focus count^10,12,13^ (Figures 1A and 1B).

**Figure 1.**
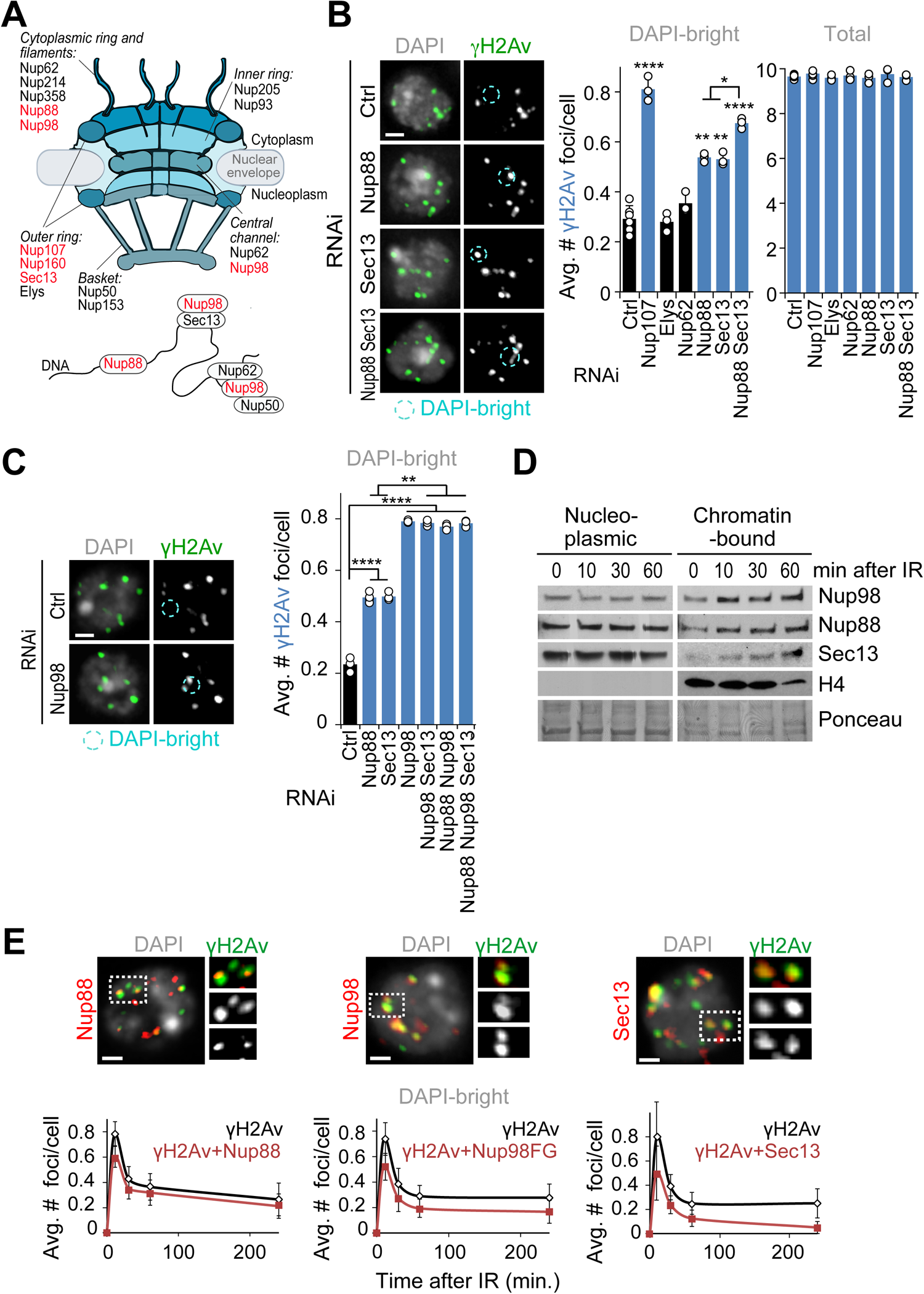
Nup88, Nup98 and Sec13 are required for relocalization of heterochromatic DSBs. (A) Schematic representation of nuclear pore proteins including chromatin-associated nucleoporins. Red indicates nucleoporins whose depletion affects relocalization/anchoring of heterochromatic DSBs from this study and^12^. (B) Immunofluorescence (IF) analysis and quantification of Kc cells fixed 60 min after IR show the number of γH2Av foci in DAPI-bright and total foci following indicated RNAi depletions. *P<0.05, **P<0.01, ****P<0.0001, n>195 cells for each RNAi condition. (C) As described in b, except cells expressing Nup96-MycFLAG were used. **P<0.01, ****P<0.0001, n>328 cells for each RNAi condition. (D) Chromatin fractionation and Western Blot (Wb) analysis of Kc cells shows the chromatin-bound fractions of Nup88, Nup98, and Sec13 at indicated time points after IR. Histone H4 and Ponceau are loading controls. (E) IF analysis shows γH2Av foci colocalizing with the indicated nucleoporins in DAPI-bright, at different timepoints after IR. Dashed boxes and zoomed details highlight colocalizations 10 min after IR. Kc cells were used for Nup88 and Sec13 staining. GFP-Nup98FG-expressing cells were used in place of Nup98 WT due to a stronger nucleoplasmic signal associated with this mutant. n>118 foci for each time point. Error bars: SEM in E, and SD in B,C from three or more independent experiments. P values calculated with two-tailed Mann– Whitney test. Images are max intensity projections of Z-stacks spanning the DAPI-bright region. Ctrl = Control. Scale bar = 1µm.

We previously showed that other nucleoporins facing the nucleoplasmic side of the pore (*i.e.,* Nup205, Nup50, Nup93 or Nup153) do not affect relocalization of repair sites^12,35^, consistent with a specialized role for the pore outer ring component Nup107 in anchoring. We tested additional nucleoporins that are known to interact with chromatin^29–31^ (Figure 1A). RNAi depletion of Nup88 or Sec13, but not RNAi of Nup62 or Elys, results in a higher number of γH2Av foci in DAPI-bright 60 min after IR relative to control RNAi (Figures 1B and S1A), consistent with relocalization defects. Nup107 and FG-porin staining at the nuclear periphery is not affected by Nup88 or Sec13 RNAi (Figures S1A and S1B), consistent with the pore being largely intact in these conditions. Notably, Nup88 and Sec13 are among the few nucleoporins identified for their nucleoplasmic function in transcription regulation, independent from their role at the pore^28–31^ (Figure 1A). Nup214 and Nup358, which closely interact with Nup88 to mediate its function at the pore^36–38^, are not required for relocalization of repair sites (Figures S1A and S1 C-E), suggesting a function of Nup88 in heterochromatin repair independent from its role at the pore.

Next, we tested the role of Nup98, another nucleoporin working both on- and off-pore (Figure 1A), which is generated from proteolysis of the Nup98-96 peptide^39^. Nup98 RNAi results in co-depletion of Nup96 and destabilization of the Y complex it belongs to, indirectly affecting Nup107 association with the pore^40^. To test the role of Nup98 independently from Nup96 and Nup107, we RNAi depleted Nup98 while exogenously expressing Nup96, which restores stability of the Y complex and Nup107 association with the nuclear pores (Figures S1A and S1B). In these conditions, Nup98 RNAi results in persistent γH2Av foci in DAPI-bright 60 min after IR, consistent with a role for Nup98 in relocalizing heterochromatic DSBs (Figures 1C and S1F).

Nup88, Nup98 and Sec13 could be required for relocalization by acting in the nucleoplasm. Accordingly, Nup88, Nup98, and Sec13 become progressively more associated with the chromatin starting 10 min after IR (Figures 1D and S1G). These nucleoporins are also enriched at repair foci in triton-extracted samples that highlight chromatin-associated proteins (Figures 1E and S1H). Co-localization with damage sites is observed starting 10 min after IR in DAPI-bright (Figures 1E and S1H), *i.e.,* before relocalization^10,12,13^. RNAi depletion of Nup88, Nup98, or Sec13 results in loss of the corresponding signals at damage foci (Figure S1I), confirming the staining specificity. Most nucleoporin-enriched repair foci are also associated with heterochromatic epigenetic marks (Figure S1J), consistent with the enrichment of Nup88, Nup98, and Sec13 at heterochromatic DSBs. We concluded that Nup88, Nup98, and Sec13 are recruited to DSBs inside the heterochromatin domain before relocalization, and they are required for this movement.

### The nucleoplasmic fractions of Nup88, Nup98 and Sec13 are required for relocalization

We directly tested the possibility that Nup88, Nup98, and Sec13 work off-pore to relocalize heterochromatic DSBs by investigating the impact of eliminating the nucleoplasmic fraction of each of these proteins to heterochromatin repair. We expressed Nup88, Nup98 or Sec13 fused to the integral transmembrane nucleoporin Ndc1, which constitutively targets proteins to the nuclear pore^41^ (Figures 2A-C). Simultaneously, we RNAi depleted endogenous Nup88, Nup98 or Sec13 using dsRNAs against their UTRs (Figures 2A-C, S1A and S2A). Eliminating the nucleoplasmic fraction of Nup88, Nup98 or Sec13 through Ndc1 fusions results in a significant defect in relocalization of heterochromatic DSBs, with γH2Av foci remaining in DAPI bright 60 min and 4h after IR (Figures 2A-C and S2B-D), while expression of the corresponding wild-type nucleoporins in the same conditions restores normal relocalization (Figures S2E-G). This supports a key role for the nucleoplasmic fraction of Nup88, Nup98 or Sec13 in relocalization of heterochromatic DSBs.

**Figure 2.**
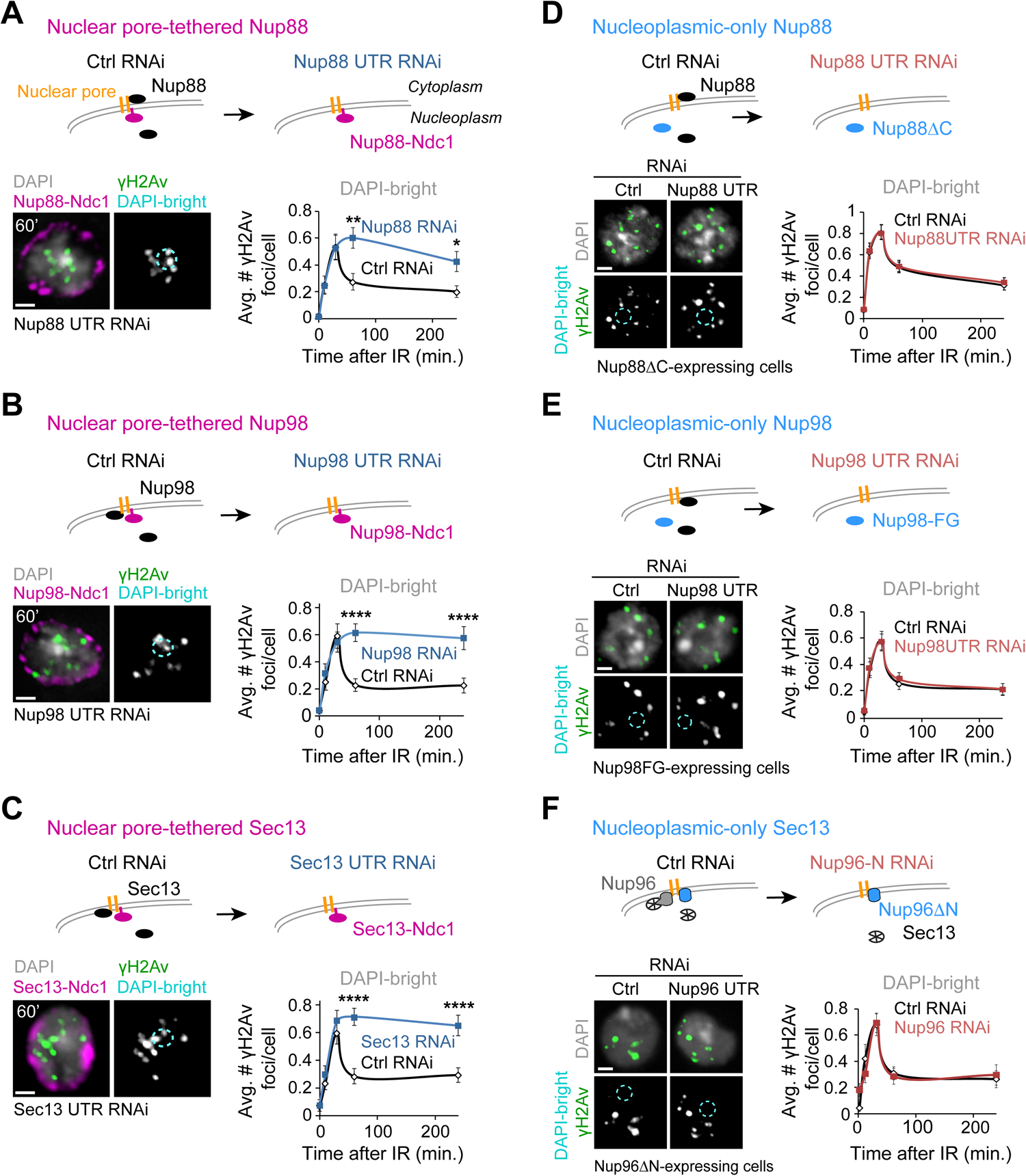
The nucleoplasmic fractions of Nup88, Nup98, and Sec13 are required for relocalization. Schematic representations (top of each figure panel) show the experimental setup. (A) Cells expressing Nup88-MycFLAG-Ndc1 are treated with dsRNAs for Nup88 UTR to deplete endogenous Nup88, or a control (Ctrl). IF (left) and quantification (right) show the number of γH2Av foci in DAPI-bright at different timepoints after IR, following indicated RNAi depletions. *P<0.05, **P<0.01, *n*>60 cells/time point. (B) As in (A), except cells express MycFLAG-Ndc1-Nup98 (plus Nup96-MycFLAG), and RNAi depletion is done against Nup98 UTR. ****P<0.0001, *n*>100 cells for each time point. (C) As in (A), except cells express Sec13-MycFLAG-Ndc1 and RNAi depletion targets Sec13 UTR. ****P<0.0001, *n*>100 cells for each time point. (D) Cells expressing FHA-Nup88ΔC are treated with dsRNAi for Nup88 UTR to deplete endogenous Nup88, or a control (Ctrl). n>77 cells for each time point. (E) As in (D), except cells express FHA-Nup98FG and dsRNAs target Nup98 UTR. *n*>100 cells for each time point. (F) As in (D), except cells express Nup96ΔN-MycFLAG and dsRNAs target only endogenous Nup96 (see also Figures S4A and S4D-F). *n*>44 cells for each time point. P values calculated with two-tailed Mann–Whitney test. Error bars are SEM. Images are max intensity projections of Z-stacks spanning the DAPI-bright region. Scale bar = 1µm.

Nup88, Nup98 or Sec13 RNAi depletions or Ndc1 fusions could affect nuclear pore transport, indirectly affecting the relocalization of heterochromatic DSBs. We tested this by measuring nucleoplasmic protein transport with the light-inducible nuclear export system LEXY^42^ (Figures S3A-B), which can detect both import and export defects^43^. Treatment with Kpt-276 or Importazole (Ipz), which affect nuclear pore export and import respectively^44,45^, results in defective transport of the LEXY construct (Figure S3C), confirming the ability of this optogenetic tool to measure nucleoplasmic transport in fly cells. Transport through the nuclear pore is largely unaffected by Nup88 or Nup98 RNAi or Ndc1 tagging, suggesting that the defects in relocalization of heterochromatic DSBs observed in these conditions are not a consequence of nuclear pore transport defects (Figures S3D-E). We detected delayed import after Sec13 RNAi and faster import in Sec13-Ndc1-expressing cells (Figure S3F). Nevertheless, we did not detect any defect in relocalization of heterochromatic DSBs after Kpt-276 or Ipz treatments (Figures S3G-H), ruling out defective pore transport as a cause of defective relocalization of heterochromatic DSBs.

As a complementary approach, we generated cells expressing “nucleoplasmic-only” versions of Nup88, Nup98 or Sec13, taking advantage of mutations that specifically disrupt anchoring of these components to the pores without affecting pore integrity^30,46,47^. As previously shown, removal of the C-terminal domain of Nup88 (Nup88ΔC) or Nup98 (Nup98FG) results in release of these proteins from the pores and loss of nuclear periphery signal^28,30,47^ (Figures S4A-C). Similarly, expression of a Nup96ΔN mutant, which loses the nuclear pore anchoring domain of Sec13^48^ (Figures S4A and S4D-E), results in Sec13 release from the nuclear periphery in the absence of endogenous Nup96 (Nup96 RNAi, Figures S4E-F). In all these conditions, recruitment of Nup88ΔC, Nup98FG, and Sec13 (in Nup96ΔN-expressing cells) to repair foci was not affected (Figure 1A for Nup98FG; Figure S4G), consistent with functional nucleoplasmic fractions of these nucleoporins. Importantly, in conditions where endogenous nucleoporins were RNAi depleted while the nucleoplasmic-only fractions of nucleoporins were expressed, relocalization of heterochromatic DSBs occurs normally, *i.e.* similarly to cells expressing endogenous nucleoporins (Figures 2D-F; S4B-C and S4F).

Together, we concluded that the nucleoplasmic fractions of Nup88, Nup98 and Sec13 are required for relocalizing heterochromatic DSBs to the nuclear periphery, and that these functions are independent from the roles of these components in nuclear pore transport.

### Sec13 and Nup88 recruit Nup98 to DSBs downstream from Smc5/6

To better understand the interplay between Nup88, Nup98, and Sec13, we investigated the epistatic relationship between these components in DSB relocalization. RNAi depletion of Nup88+Sec13 results in a higher number of γH2Av foci in DAPI-bright 60 min after IR relative to individual RNAi depletions, indicating that these two proteins independently contribute to relocalizing heterochromatic DSBs (Figure 1B). Interestingly, Nup88+Nup98, Nup98+Sec13, and Nup88+Nup98+Sec13 RNAi lead to relocalization defects similar to Nup98 depletion alone (Figures 1C and S1F), consistent with Nup98 working with both Nup88 and Sec13 for relocalization. In addition, RNAi depletion of Nup88 or Sec13 significantly lowers the number of Nup98 foci colocalizing with γH2Av (Figure 3A), revealing a role for Nup88 and Sec13 in Nup98 loading to DSBs, similar to the role of Sec13 in Nup98 recruitment to transcription sites^29^. Both Nup88 and Sec13 are β-propeller proteins^49^, suggesting the importance of this structure in Nup98 recruitment to damage sites.

**Figure 3.**
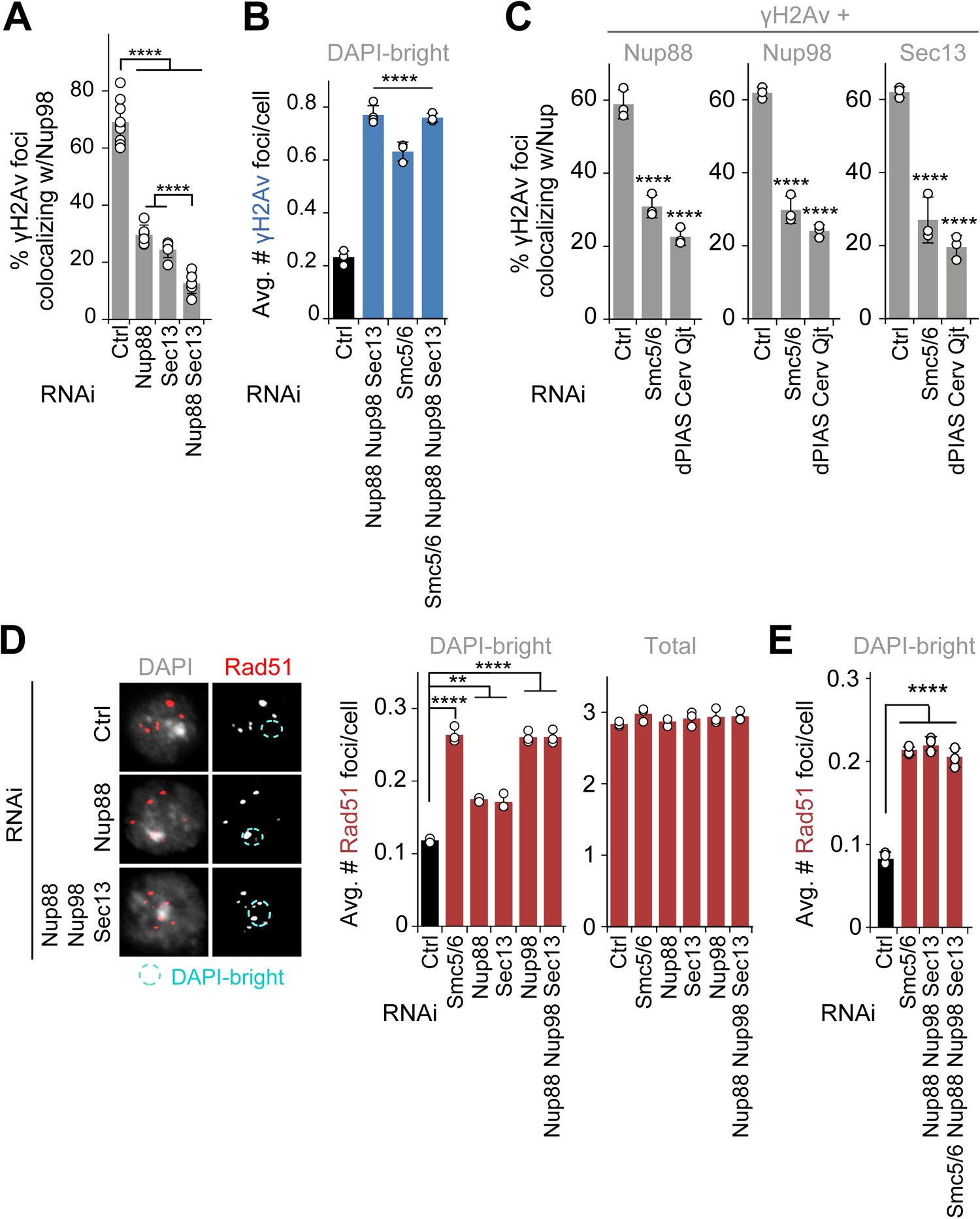
Nup88, Nup98 and Sec13 act downstream from Smc5/6 to promote relocalization and to prevent abnormal Rad51 recruitment. (A) Quantification of cells fixed 10 min after IR shows the number of γH2Av foci colocalizing with GFP-Nup98FG foci in DAPI-bright following indicated RNAi depletions. ****P<0.0001, *n*>74 foci for RNAi condition. (B) Quantification of cells expressing Nup96-MycFLAG fixed 60 min after IR, shows the number of γH2Av foci in DAPI-bright after indicated RNAi depletions. ****P<0.0001 relative to Ctrl, n>226 cells for each condition. (C) Quantification of cells expressing Nup88-MycFLAG, GFP-Nup98FG or Sec13-MycFLAG shows colocalizations with γH2Av foci 10 min after IR, following indicated RNAi depletions. ****P<0.0001 relative to Ctrl, n>495 foci for each condition. (D) IF and quantification of cells fixed 60 min after IR show the number of Rad51 foci in DAPI-bright and total foci following indicated RNAi depletions. Smc5/6 RNAi results in abnormal Rad51 focus formation in heterochromatin^10,12,15^ and is used as positive control. n>246 cells for each condition. **P<0.01, ****P<0.0001 relative to Ctrl. (E) As in (D). ****P<0.0001 for comparisons to Ctrl, n>581 cells for each RNAi condition. Error bars are SD from three independent experiments. P values calculated with two-tailed Mann–Whitney test. Scale bar = 1µm.

We further investigated the mechanisms responsible for Nup88-Nup98 and Sec13-Nup98 recruitment to repair sites. Previous studies showed that Nup98 recruitment to transcription sites requires the histone modifying complex MBD-R2^50^. Human Nup98 also interacts with the DExH-Box Helicase 9 (Dhx9)^51^, which contributes to HR repair^52,53^. However, RNAi depletion of MBD-R2 or Dhx9 does not affect relocalization of heterochromatic DSBs (Figures S5A and S5B), ruling out a role for these components in recruiting nucleoporins to repair sites for relocalization.

We turned our attention to Smc5/6, which is required for relocalization of heterochromatic DSBs and acts downstream from silencing chromatin marks and resection in this function^10,12,15^. RNAi depletion of Nup88+Nup98+Sec13+Smc5/6 results in similar relocalization defects as Nup88+Nup98+Sec13 RNAi (Figures 3B and S5C), revealing that Nup88, Nup98 and Sec13 work in the same pathway as Smc5/6 for relocalization.

We investigated which steps of the Smc5/6-mediated relocalization pathway are dependent on Nup88, Nup98 and Sec13, by testing how depletion of these nucleoporins affects: damage signaling and resection (*i.e*., Mu2/Mdc1 or ATRIP foci^10^), Arp2/3 and myosin recruitment to repair sites^13^, and Nse2 recruitment to heterochromatin and repair sites^10,12^. Nucleoporin RNAi does not affect Mu2/Mdc1 and ATRIP focus formation (Figure S5D); Arp2/3, Myosins or Unc45 recruitment to DSBs (Figure S5E); or Nse2 recruitment to heterochromatin and repair foci (Figure S5D). Consistent with normal Arp2/3 recruitment, nuclear actin filaments also form normally in the absence of nucleoporins (Figure S5F). On the other hand, RNAi depletion of Smc5/6 or SUMO E3-ligases (e.g., dPIAS, Nse2/Cerv and Nse2/Qjt) affects Nup88, Nup98 and Sec13 enrichments at γH2Av foci (Figure 3C), placing nucleoporins recruitment to repair sites downstream from Smc5/6 and SUMOylation.

In addition to mediating relocalization, Smc5/6 is required to exclude Rad51 foci from the heterochromatin domain^6,10,12^. Thus, we tested the role of nucleoporins in this function. Nup88, Nup98 or Sec13 RNAi results in abnormal formation of Rad51 foci inside the heterochromatin domain 60 min after IR (Figures 3D and S5G), without affecting the total number of Rad51 foci (Figures 3D and S5G). Removal of Nup98 or all three nucleoporins together results in a higher number of Rad51 foci specifically in DAPI-bright, relative to the individual RNAi depletion of Nup 88 or Sec 13 ( Figures 3 D and S 5 G). Additionally, RNAi depletion of Smc5/6+Nup88+Nup98+Sec13 results in a similar number of Rad51 foci inside the heterochromatin domain, relative to the RNAi depletion of Smc5/6 alone (Figures 3E and S5H), indicating that nucleoporins mediate Rad51 exclusion in the same pathway as Smc5/6.

Together, these results revealed that Sec13 and Nup88 independently recruit Nup98 to repair sites, and that these components act downstream from Smc5/6 for relocalization of heterochromatic DSBs and for Rad51 focus exclusion from the domain.

### Nucleoporins are required for mobilizing and clustering heterochromatic DSBs

The nucleoplasmic fractions of Nup88, Nup98 and Sec13 could promote relocalization of heterochromatic DSBs either by facilitating the mobilization of repair foci, or by tethering repair sites to the nuclear periphery after relocalization similar to Nup107. We investigated this directly by characterizing the dynamics of repair sites using live imaging of Mu2/Mdc1 foci originating from the heterochromatin domain marked by mCh-HP1a^10,12,13,54^.

We previously showed that RNAi depletion of anchoring components at the nuclear periphery results in a higher plateau of the Mean Square Displacement (MSD) curve describing the dynamics of heterochromatic repair foci, consistent with increased exploration of the nuclear space^12^. In striking contrast, RNAi depletion of Nup88, Nup98 and Sec13 leads to a drastic reduction of the plateau of the MSD curve, reflecting decreased mobility (Figure 4A). Consistent with reduced dynamics, repair foci remain inside the heterochromatin domain throughout the 1h time course in the absence of Nup88, Nup98 and Sec13, while foci typically reach the nuclear periphery during the same timeframe in control conditions (Figures 4A-C). These results are consistent with a role for Nup88, Nup98 and Sec13 in mobilizing heterochromatic DSBs, rather than in anchoring repair sites to the nuclear periphery. Remarkably, the dynamics of euchromatic repair foci are largely unaffected by Nup88, Nup98 and Sec13 RNAi (Figure 4A), revealing a role for these nucleoporins in mobilizing repair sites specifically in heterochromatin.

**Figure 4.**
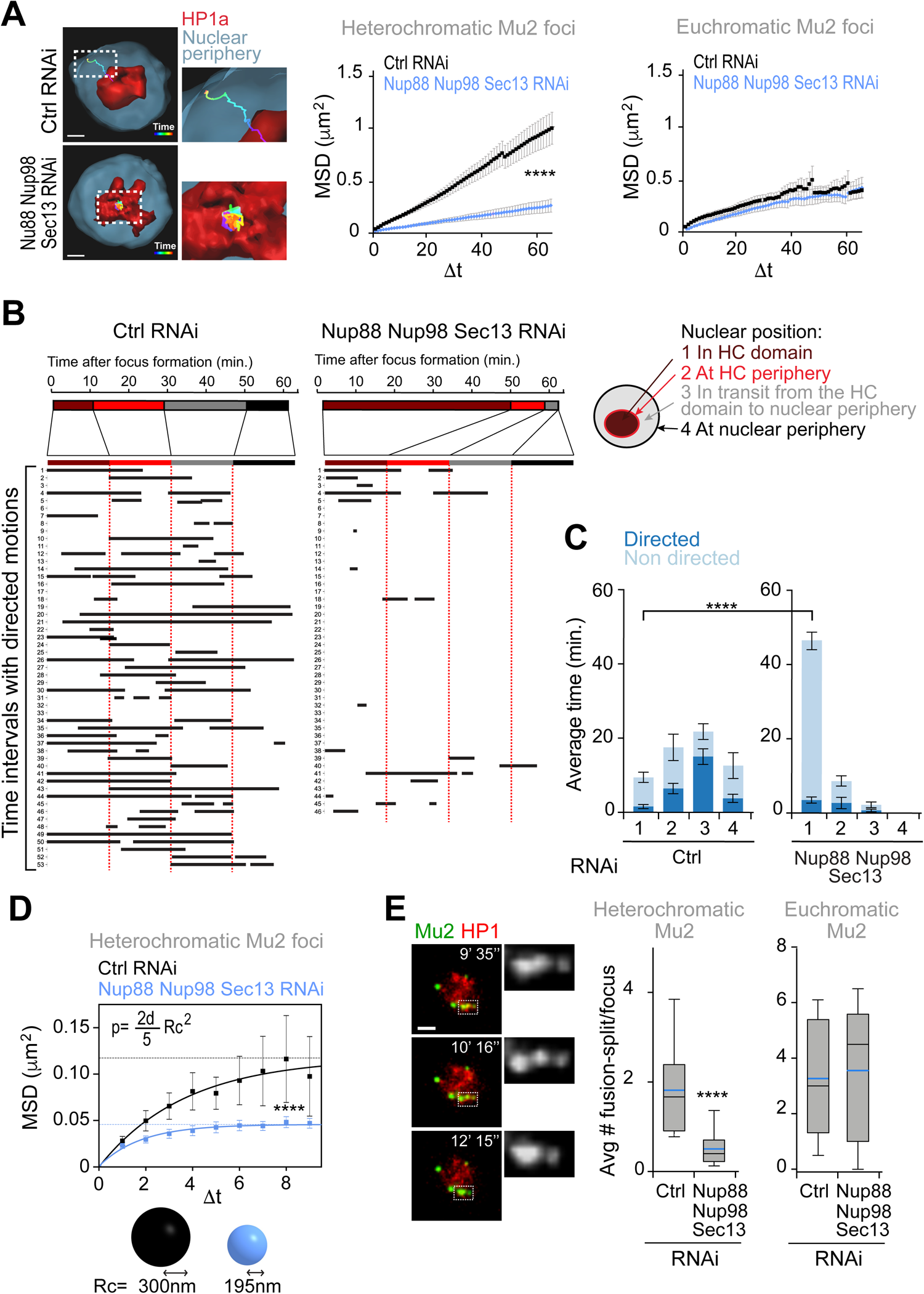
Nup88, Nup98, and Sec13 nucleoporins are required for mobilizing heterochromatic DSBs. (A) 3D reconstructions in Imaris (left) and MSD analysis (right) of GFP-Mu2 foci that form inside (heterochromatic) or outside (euchromatic) the mCh-HP1a domain after IR, following indicated RNAi depletions. n≥46 tracks for each condition from three or more experiments. Dashed boxes and zoomed details highlights the tracks. Δt, time intervals (40s each). ****P>0.0001. (B) LDM analysis of heterochromatic foci from the experiment described in (A). Duration and distribution of LDMs (black lines) along each track relative to different nuclear positions, adjusted relative to a ‘pseudo-trajectory’ defined by the average time that foci spend in each nuclear space, and displayed for equalized time intervals. Numbers on the left side indicate individual foci. (C) Average duration of directed and non-directed motions for heterochromatic foci in (A,B), in the indicated nuclear locations. ****P>0.0001. (d) MSD analysis of time points from (A) characterized by non-directed motion inside the heterochromatin domain, following indicated RNAi depletions. ****P>0.0001, n=29 foci for Ctrl and n=38 foci for Nup88+Nup98+Sec13 RNAi from (A). The confinement radius (below, Rc) was calculated using the plateau of the curve (p, dashed line), as indicated in^54^. d: dimensions (d=3 in this analysis) (E) Images from a movie (left) and quantification (right) show fusion and spitting events for heterochromatic Mu2/Mdc1 foci in cells expressing GFP-Mu2 and mCh-HP1a, following indicated RNAi depletions. Dashed boxes and zoomed details highlight fusion-splitting events. In the box plot blue line is the mean; black line is the median n>97 foci per each condition. ***P<0.001 for the indicated comparison. Error bars are SEM in A, C, D. The whiskers are drawn down to the 10th percentile and up to the 90th in e. P values calculated with extra sum-of-squares F-test, nonlinear regression for curve fitting for A, D and two-tailed Mann–Whitney test in C, E. Scale bars = 1 µm.

Next, we investigated which component of the movement is most affected in the absence of Nup88, Nup98 and Sec13 by identifying the time points associated with directed or subdiffusive motion. We previously detected long directed motions (LDMs) associated with repair foci that leave the heterochromatin domain, which mostly occur in transit from the heterochromatin domain periphery to the nuclear periphery, *i.e.,* where most nuclear actin filaments are assembled^9,13,54^ (Figures 4B-C and S6A,B). Inside the heterochromatin a significant portion of the movement is subdiffusive, in agreement with the existence of constraints on the movement and the lack of actin filaments^9,13^. RNAi depletion of Nup88, Nup98 and Sec13 results in a significant reduction of LDM frequency and duration, in agreement with most repair sites remaining inside the heterochromatin domain (Figures 4B-C and S6A-B). However, the few repair sites that leave the domain still display directed motion, suggesting that LDMs are not primarily affected once repair foci are released from the heterochromatin domain (Figures 4B and S6A-B). This is consistent with normal Arp2/3 and myosin recruitment to repair sites as well as normal actin filament formation following RNAi depletion of Nup88, Nup98 and Sec13 (Figures S5E-F).

Intriguingly, MSD analysis of the time points characterized by subdiffusive motion inside the heterochromatin domain reveals a significant reduction in the space explored during this initial phase of movement in the absence of Nup88, Nup98 and Sec13 (Figure 4D). This suggests that the primary consequence of losing these nucleoporins is a reduced overall mobility inside the heterochromatin domain, which in turn affects the ability of foci to reach the heterochromatin domain periphery and initiate directed motions. We also detected a reduced ability of repair foci to fuse and split inside the heterochromatin domain upon RNAi depletion of Nup88, Nup98 and Sec13, consistent with reduced dynamics (Figure 4E). Consistent with a primary role for the nucleoporins in heterochromatin repair, their RNAi depletion does not affect the frequency of focus fusion and splitting events in euchromatin (Figure 4E).

We conclude that Nup88, Nup98 and Sec13 nucleoporins mobilize repair sites inside the heterochromatin domain, promoting fusion and fission events and facilitating focus relocalization to the periphery of the heterochromatin domain independently from F-actin and myosins.

### Nup98 phase separation properties are required and sufficient for relocalization of heterochromatic repair sites and Rad51 exclusion

The Nup98FG truncated protein that is able to carry out relocalization of heterochromatic DSBs in the absence of endogenous Nup98 (Figures 2E, S4A and S4C) is mostly composed of phenylalanine-glycine (FG) repeats^30^ (Figure S4A). These “FG-domains” are common components of nucleoporins that face the nuclear pore channel, where they create a phase separated environment *via* hydrophobic and aromatic (π-π) interactions between phenylalanine residues^24–27^. Consistently, Nup98 and the Nup98FG domain spontaneously phase separate *in vitro* at physiologic concentrations^24,27^. This property is highly conserved across species, including in *Drosophila*^27^. In agreement with their ability to phase separate, Nup98 FG-domains mostly consist of intrinsically disordered regions (IDRs) (Figure S7A). Phase separation properties are also consistent with the role of Nup98 in repair focus fusion and splitting behavior (Figure 4E). Thus, we tested whether the phase separation properties of Nup98FG are required for relocalization of heterochromatic DSBs.

We generated mutant forms of Nup98FG that are either unable to phase separate (Nup98AG) or that restore hydrophobic interactions and phase separation properties (Nup98YG) by mutating all the FG repeats into AG or YG, respectively^24,26^ (Figures 5A and S4A). We confirmed the effects of these mutations *in vivo* using an optoDroplet assay, where IDRs are fused to photoreceptor cryptochrome 2 (Cry2) of *A. thaliana* to promote self-association upon blue light exposure^20,55,56^ (Figure S7B). When activated by light stimulation, Cry2-mCherry-Nup98FG and Cry2-mCherry-Nup98YG rapidly form droplets (Figures S7C-D and Supplementary movie S1), which undergo fusion and fission (Figure S7C), confirming their phase separation properties. Furthermore, these droplets are quickly reversed upon light removal and can be re-induced by light exposure in the same cell multiple times (Figures S7E-F). Conversely, Cry2-mCherry-Nup98AG does not form droplets, similar to the negative control Cry2-mCherry, confirming its inability to phase separate (Figures S7C-D).

**Figure 5.**
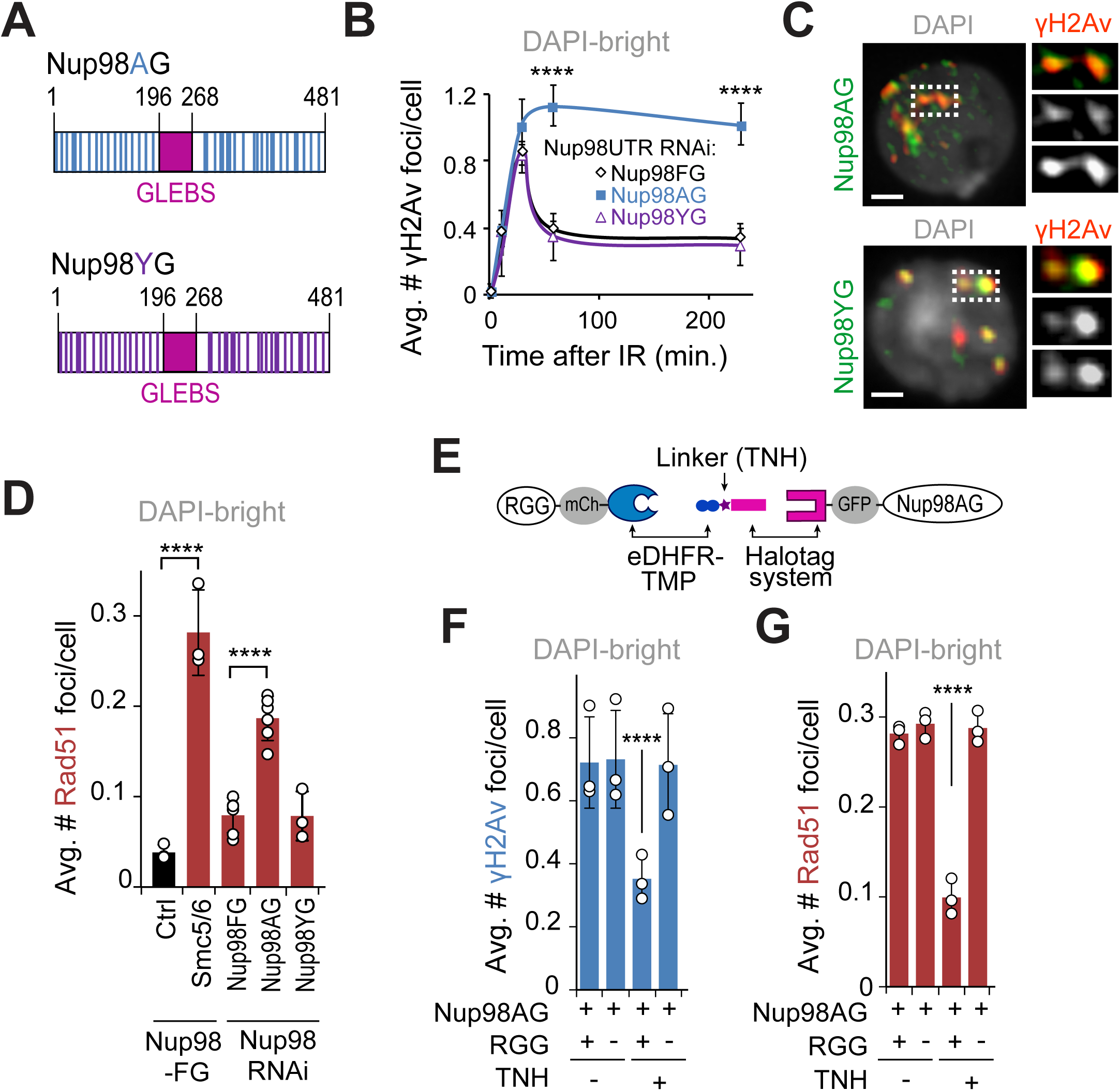
Nup98 mediates relocalization of heterochromatic DSBs *via* phase separation. (A) Schematic representation of Nup98AG and YG mutations. Each Phenylalanine of the FG-domain of Nup98FG is substituted with Ala in Nup98AG or Tyr in Nup98YG. (B) Quantification of γH2Av foci in DAPI-bright at different time points after IR, in cells expressing FHA-Nup98-FG, -AG or -YG following RNAi depletion of endogenous of Nup98 (Nup98 UTR RNAi). \*\*\*\**P*<0.0001 for Nup98AG relative to Nup98FG, n>50 cells for each time point. (C) IF shows Nup98AG and Nup98YG foci colocalizing with γH2Av foci. Dashed boxes and zoomed details highlight colocalizations. (D) Quantification of cells expressing FHA-Nup98-FG, -AG or -YG following depletion of endogenous Nup98, fixed 60 min after IR, shows the number of Rad51 foci in DAPI-bright. \*\*\*\**P*<0.0001 for indicated comparisons. *n*>363 cells for each RNAi condition. Smc5/6 RNAi in cells expressing Nup98FG is shown as a positive control. (E) Schematic representation of the dimerizer system (adapted from^58^). The RGG domains are tagged with mCherry (mCh) and fused to eDHFR, whereas Nup98AG is tagged with GFP and fused to 3xHalo. The dimerizer is TNH. (F) Quantification of cells expressing (+) or non-expressing (-) RGG-mCherry-RGG-eDHFR, plus Halo-GFP-Nup98AG, treated for 120 min with (+) or without (-) TNH as indicated, and fixed 60 min after IR, shows the number of γH2Av foci in DAPI-bright. ****P<0.0001. n>346 cells for each condition. (G) As in F, except Rad51 foci were quantified. ****P<0.0001. n>342 cells for each condition. Error bars are SEM in B, and SD in D,F,G from three independent experiments. P values calculated with two-tailed Mann–Whitney test. Scale bars = 1 µm.

Next, we tested the effect of these mutations on relocalization of heterochromatic repair foci. Similar to Nup98FG, expression of Nup98YG in the absence of endogenous Nup98 results in normal relocalization of repair foci (Figures 5B and S8A). Conversely, expression of the phase separation mutant Nup98AG results in heterochromatic breaks remaining inside the heterochromatin domain 60 min and 4h after IR, consistent with a major relocalization defect (Figures 5B and S8A). Notably, Nup98AG and Nup98YG are enriched at repair foci similarly to Nup98FG (Figures 1E and 5C), establishing that defective relocalization is not a consequence of defective recruitment of Nup98AG to damage sites.

A typical property of phase separated structures is selective permeability, *i.e.,* the ability to exclude certain proteins while enabling the passage of others. Thus, we tested the ability of Nup98-mediated phase separation to exclude Rad51 from repair foci inside the heterochromatin domain. Remarkably, expression of Nup98AG in the absence of endogenous Nup98 results in abnormal formation of Rad51 foci inside the heterochromatin domain (Figures 5D and S8B,C). Nup98YG expression restores Rad51 exclusion from the domain, similar to Nup98FG (Figure s5D and S8B,C), revealing the importance of Nup98-mediated condensates in Rad51 exclusion from repair sites inside the heterochromatin domain.

Relocalization of heterochromatic γH2Av foci and exclusion of Rad51 are also defective after cell treatment with 1,6-Hexanediol (Figures S8D,E), an aliphatic alcohol that weakens hydrophobic interactions and interferes with phase separation^26,57^, consistent with the importance of phase separation in these functions.

We next tested whether Nup98-dependent phase separation is sufficient to induce repair focus relocalization and Rad51 exclusion in heterochromatin by investigating whether artificially inducing condensate formation is sufficient to rescue Nup98AG mutant defects. To achieve this, we used a dimerization system that fuses Nup98AG with two RGG domains that induce condensates in the presence of the chemical linker TMP-NVOC-Halo (TNH)^58,59^ (Figure 5E). Addition of TNH to the cell media results in the formation of Nup98AG condensates (Figure S8F). Strikingly, this is sufficient to restore normal relocalization of heterochromatic repair sites and Rad51 exclusion from the heterochromatin domain (Figures 5F,G and S8G,H).

Together, these studies revealed that the phase separation properties of Nup98 are required and sufficient to relocalize heterochromatic repair foci and to block Rad51 recruitment inside the heterochromatin domain.

### Nup98 phase separation properties are required for heterochromatin repair and stability

These studies identified nucleoplasmic Nup88, Nup98 and Sec13 as new components required for relocalization of heterochromatic DSBs and for preventing Rad51 recruitment inside the heterochromatin domain, suggesting the importance of these proteins in the progression of heterochromatin repair. We directly tested this by determining the biological consequences of perturbing these nucleoporins and the phase separation properties of Nup98 on IR sensitivity, damage processing, and genome stability.

First, RNAi depletion of Nup88 or Sec13 (Figure S9A) or removal of the nucleoplasmic fraction of nucleoporins through Ndc1-tagging (Figure 6A) results in IR sensitivity, consistent with defective damage repair^12,13^. Conversely, expression of the nucleoplasmic-only fraction of the nucleoporins (Nup88ΔC, Nup98FG, or Sec13 in a Nup96ΔN background, respectively) results in IR sensitivity comparable with controls expressing wild-type nucleoproteins (Figure 6A). Similarly, expression of Nup98AG mutants in the absence of endogenous Nup98 results in IR sensitivity, which is rescued by expression of Nup98YG (Figure 6B). This is consistent with the importance of the nucleoplasmic fraction of Nup88, Nup98 and Sec13, and particularly the phase separation properties of Nup98, in IR resistance.

**Figure 6.**
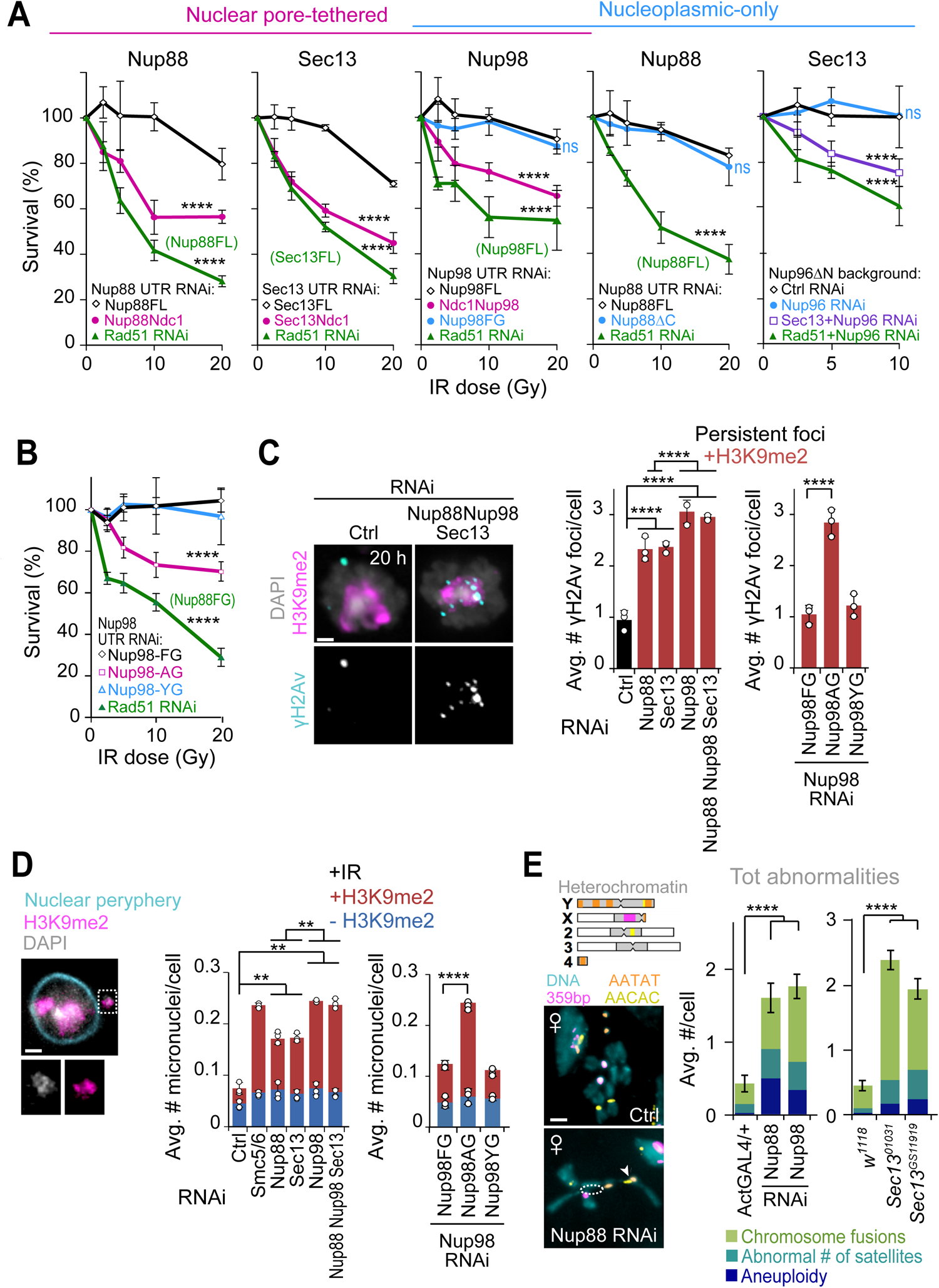
Nup88, Nup98, and Sec13 nucleoporins are required for heterochromatin repair and stability. (A) IR sensitivity assays show cell survival at different IR doses and after indicated RNAi depletions and protein expression. Rad51 RNAi results in cell sensitivity to IR^12,13^ and is used as positive control in cells expressing each FL (full-length) protein or Nup96ΔN, as indicated. (Left to right) First graph: Survival of cells expressing Nup88-MycFLAG-Ndc1 or Nup88FL-MycFLAG in the absence of endogenous Nup88 (Nup88 UTR RNAi). ****P<0.0001 for all comparisons *vs* Nup88FL. Second graph: Survival of cells expressing Sec13-MycFLAG-Ndc1 or Sec13FL-MycFLAG, after Sec13 UTR RNAi. ****P<0.0001 for comparisons *vs* Sec13FL. Third graph: Survival of cells expressing FHA-Nup98FL, MycFLAG-Ndc1-Nup98, or FHA-Nup98FG, after Nup98 UTR RNAi. ****P<0.0001 for Ndc1-Nup98 or Rad51 RNAi *vs* Nup98FL, P=ns for Nup98FG *vs* Nup98FL. Fourth graph: Survival of cells expressing FHA-Nup88ΔC or FHA-Nup88FL, after Nup88 UTR RNAi. P=ns for Nup88ΔC vs Nup88FL. ****P<0.0001 for Rad51 *vs* FHA-Nup88FL. Fifth graph: Survival of cells expressing Nup96ΔN-MycFLAG after RNAi depletion of: Ctrl; endogenous Nup96 (Nup96 UTR RNAi); Sec13+Nup96 (positive control); or Rad51+Nup96 (positive control). ****P<0.0001 for comparisons *vs* Ctrl RNAi, P=ns for Nup96 vs Ctrl RNAi. n>150 cells per RNAi per dose. (B) As in (A), except cells expressing FHA-Nup98-FG, -AG or -YG were used, following Nup98 UTR RNAi depletion. ****P<0.0001 for Nup98AG or Rad51 *vs* Nup98FG. n>100 cells per RNAi per dose. Rad51 RNAi is used as positive control in cells expressing Nup98FG. (C) IF and quantifications of cells fixed 20 hrs after IR and stained for γH2Av and H3K9me2 following indicated RNAi depletions, or in cells expressing FHA-Nup98-FG, -AG or -YG following Nup98 UTR RNAi. ****P<0.0001, n>273 cells for each condition. +H3K9me2: γH2Av foci associated with H3K9me2 signals. (D) IF and quantification of micronuclei in cells stained for DNA (DAPI), heterochromatin (H3K9me2), and nuclear periphery (Nup62), after indicated RNAi depletions or in cells expressing FHA-Nup98-FG, -AG or -YG, after Nup98 UTR RNAi depletion, 2 days post IR. **P<0.01, ****P=0.0001, n>447 cells for each condition. +/− H3K9me2: micronuclei with/ without H3K9me2. Dashed box and the zoomed detail highlights a micronucleus. (E) Images of chromosome preparations from larval brains and quantifications from Ctrl and indicated RNAi depleted or mutant larval brains stained by FISH for indicated satellites. Examples of fusion (arrowhead) and chromosome arm losses (dashed circle) are highlighted. n>46 karyotypes/ genotype from at least two independent crosses and three brains. Nup98 RNAi lines also express Nup96-Myc^77^. ****P<0.0001 for indicated comparisons. The diagram of *Drosophila* chromosomes indicates satellite positions. Error bars are SEM in E and SD from three or more experiments in A-D. P values calculated with extra sum-of-squares *F*-test, nonlinear regression for curve fitting for A and B, two-tailed Mann–Whitney test in C-D, and unpaired T-test in E. Images are max intensity projections of a few Z-stacks. Scale bar = 1µm.

Second, RNAi depletion of Nup88, Nup98 or Sec13 results in persistent γH2Av foci associated with the heterochromatin domain 20h after IR (Figures 6C and S9B), a time point at which heterochromatin repair is largely completed in control RNAi conditions or in euchromatin (Figures 6C and S9B)^12,13^. Stronger defects are observed after RNAi depletion of Nup98 or the simultaneous depletion of all three nucleoporins (Figure 6C), consistent with Nup88-Nup98 and Sec13-Nup98 working independently for heterochromatin repair (Figures 1B,C and 3A). Similar persistent foci are detected when cells express the phase separation mutant construct Nup98AG, but not upon expression of Nup98YG, in agreement with with the importance of Nup98-driven phase separation in resolving heterochromatic DSBs (Figures 6C and S9B).

Third, RNAi depletion of Nup88, Nup98 or Sec13 triggers the formation of damage-induced heterochromatic micronuclei, with the strongest phenotype observed after Nup98 or Nup88+Nup98+Sec13 RNAi (Figures 6D and S9C). This is mimicked by expression of Nup98AG and not by expression of Nup98YG (Figures 6D and S9C), establishing the importance of Nup98-mediated phase separation in preventing micronucleus formation.

RNAi depletion or mutation of Nup88, Nup98 or Sec13 also results in genome instability in *Drosophila* larval neuroblasts (Figures 6E and S9D-F), where abnormal karyotypes likely originate due to defective repair of spontaneous DSBs during larval development. Chromosome rearrangements include aneuploidies, chromosome fusions and changes in the number of satellites (Figures 6E and S9F), most of which involve the heterochromatic fourth and Y chromosomes or pericentromeric regions, as expected for defective heterochromatin repair (Figures 6E and S9F)^12,13^. Similar to experiments in cultured cells, RNAi depletion of Nup98 was done in flies expressing an exogenous copy of Nup96-Myc to avoid damages to the outer ring of the nuclear pore through Nup96 depletion^50^. Accordingly, Nup107 and MAb414 staining at the nuclear periphery is not affected after Nup98 RNAi in these flies (Figure S9E). Together, we conclude that Nup88, Nup98 and Sec13 nucleoporins, and particularly the phase separation property of Nup98, are critical for heterochromatin repair and stability in *Drosophila* cells and tissues.

## DISCUSSION

This study revealed a novel role for nucleoporins in heterochromatin repair and genome stability. Nup88 and Sec13 independently recruit Nup98 to heterochromatic DSBs downstream from Smc5/6, and these components are required to relocalize repair sites and prevent Rad51 recruitment inside the heterochromatin domain. The nucleoplasmic function of these nucleoporins, and not their association with the nuclear pore, is specifically required for relocalization, revealing a novel off-pore role for Nup88, Nup98 and Sec13 in DNA repair. Importantly, these nucleoporins mediate the mobilization of repair sites, promoting their clustering and subdiffusive motion at early relocalization steps, and this function is specific for heterochromatic DSBs. Nucleoporin-induced motion precedes and is independent from the nuclear F-actin and myosin-driven directed motions that guide repair sites from the periphery of the heterochromatin domain to the nuclear periphery. The phase separating properties of Nup98 are required and sufficient to promote DSB mobilization while also suppressing Rad51 recruitment, thus preventing aberrant recombination. Consistently, misregulation of this pathway results in heterochromatin repair defects and instability in cells and tissues. Together, these results support a model in which the nucleoplasmic fraction of Nup98 induces phase separation to create a selective barrier that blocks Rad51 access to heterochromatic repair foci, while promoting focus clustering and the diffusive motion of repair foci to the heterochromatin domain periphery. Once at the domain periphery, repair foci initiate directed motion to the nuclear periphery along actin filaments for HR repair progression (Figure 7).

**Figure 7.**
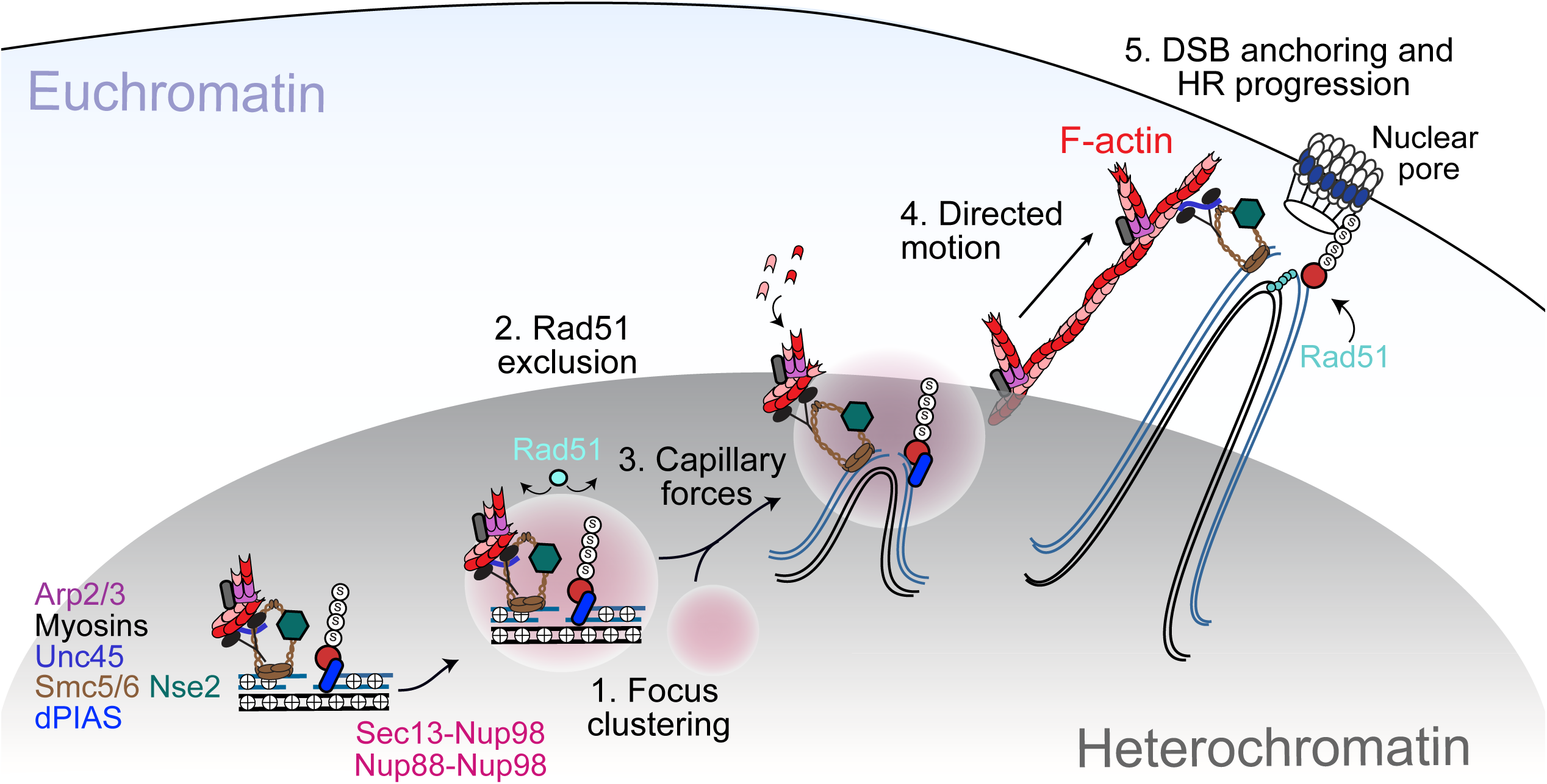
Model for the role of Nup88, Nup98 and Sec13 in heterochromatic DSBs repair in *Drosophila*. DSBs are resected inside the heterochromatin domain, where Smc5/6 promotes the loading of Nup88-Nup98 and Nup98-Sec13. Nup98-FG repeats create a phase separated environment that facilitates focus clustering inside the heterochromatin domain while excluding Rad51. We hypothesize that a multiphase condensate established by Nup98 inside the HP1a domain generates capillary forces that promote the relocalization of Nup98 condensates at repair sites to the surface of the heterochromatin domain. Here, Nup98-driven wetting behavior might also promote the association of repair foci with actin filaments generated by Arp2/3. Unc45 recruitment by Smc5/6 activates nuclear myosins to “walk” repair sites along filaments, thus relocalizing DSBs to nuclear pores where HR continues with Rad51 recruitment. Relocalization prevents ectopic recombination by isolating damaged sites and their homologous templates (black lines) from undamaged repeats before strand invasion.

How nucleoporins promote the mobilization of heterochromatic DSBs independently from nuclear F-actin and myosins is intriguing. Nucleoporins are required for focus fusion/splitting and diffusion inside the heterochromatin domain, and this is consistent with repair foci and heterochromatin behaving as phase separated structures. We propose that the initial mobilization of heterochromatic repair sites relies on the establishment of a multiphase condensate, with Nup98-dependent condensates forming inside the phase separated heterochromatin domain^22,23^. The formation of a condensate inside another condensate, in conditions where the two liquid phases are immiscible, results in capillary forces due to interfacial tension, which push one condensate to the surface of the other^60,61^. These forces alone could explain how Nup98-containing condensates promote relocalization of DSBs to the surface of the heterochromatin domain.

It is also noteworthy that fusing RGG domains to the Nup98AG phase separation mutant can rescue relocalization defects and the exclusion of Rad51 during the damage response. This suggests that the biophysical properties established by RGG domains mimic some or all of the properties established by FG repeats during repair. Thus, the biophysical properties are likely more important than the sequence composition of the seeding molecules for establishing the functions of Nup98 condensates in heterochromatin repair.

Collectively, our study suggests that Nup98-dependent phase separation contributes to heterochromatin repair at multiple levels (Figure 7): i) by promoting repair focus clustering, which likely promotes early repair steps by increasing the local concentration of repair components; ii) by facilitating the release of repair foci from the heterochromatin domain through capillary forces; and iii) by excluding Rad51 from heterochromatic repair foci through selective permeability, thus preventing aberrant recombination between repeated sequences.

The identification of Nup98 as a new DSB repair component working through phase separation also raises new questions about how these structures interface with other condensates established by repair proteins at DSBs^62^, such as those formed by MRNIP^63^, PARP^64^, Rap80^65^, RPA^66^, TopBP1^67^, Rad52^68,69^, Slx4^70^, FUS^63,64,71^ and 53BP1^56,72^. Phase separated proteins can promote growth and changes in properties of pre-existing condensates through miscible properties^61^, or maintain immiscible condensates with reduced protein exchanges^21^, each specialized in a distinct repair step^21^. Consistent with this idea, it has been proposed that the interface between immiscible condensates allows for protein exchange, facilitating sequential biological reactions^21^. It is also tempting to speculate that the interaction between phase separated repair foci and nuclear F-actin affects nuclear dynamics^73^. For example, the wetting properties of phase separated structures could facilitate interaction and sliding of repair foci along F-actin^68,73^. Understanding the properties of and the relationships between these structures is therefore an important future goal to better understanding the impact of condensates on repair focus dynamics and repair progression.

Importantly, our studies also suggest new mechanisms for nucleoporin-mediated oncogenic transformation. All Nup98-associated leukemias contain the Nup98FG domain fused with transcription factors, chromatin modifiers, or other chromatin-associated proteins^74^, and the oncogenic potential of Nup98 has been ascribed to its ability to form condensates^32–34^. The current model proposes that bringing the FG domain to transcription sites promotes myeloid transformation by altering gene expression^34,74^. Our study suggests alternative mechanisms for oncogenic transformation: Nup98 oncofusions can titrate Nup98 away from repair sites or abnormally recruit transcriptional activators to heterochromatic repair sites, either of which would hijack the repair response in heterochromatin, inducing massive genomic instability. Similarly, higher levels of Nup88 have been linked to higher cancer aggressiveness^75,76^, which could also derive from misregulation of heterochromatin repair.

Together, these studies establish a new role for nucleoplasmic nucleoporins in nuclear dynamics, heterochromatin repair, and genome stability in a multicellular eukaryote, providing a new framework for understanding how their misregulation can contribute to cancer initiation and progression.

## Supplemenatry Figure Legends

**Supplementary Figure 1. Relatated to figure 1. Nup88, Nup98 and Sec13 are required for relocalization of heterochromatic DSBs** (A) Wb analyses show Sec13, Elys, and Nup98 RNAi depletion efficiencies. Lamin, Actin or (*) a non specific band, are used as loading controls. qPCR analyses show Nup62, Nup88 and Nup358 RNAi depletion efficiencies. GFP fluorescence intensity normalized for the GFP background shows Nup98 RNAi efficiency in GFP-Nup98FG-expressing cells. ****P<0.0001, n>20 cells. (B) Top: IF analysis shows that RNAi depletions of Nup88 or Sec13 does not affect Nup107 signal at the nuclear periphery or FG-porin staining with MAb414 antibodies. Bottom: IF analysis shows that Nup98 RNAi depletion does not affect Nup153 localization. Given that loss of Nup107 from the nuclear periphery affects Nup153 localization^12^, this also indicates normal Nup107 association with pores. (C) Wb and IF analyses (top) show that Nup214 RNAi depletion affects Nup88 protein level, as previously shown^36^. Wb analysis (bottom) shows that exogenous expression of Nup88ΔC is not not affected by Nup214 RNAi. (D-E) Quantification of cells fixed 60 min after IR shows the number of γH2Av foci in DAPI-bright and total foci following indicated RNAi depletions, n>218 cells for each RNAi condition. **P<0.001; ***P<0.001. For Nup214 RNAi experiments, cells expressing Nup88ΔC were used to rescue Nup88 levels as shown in C. (F) Quantification of the experiment described in Figure 1C shows the total number of γH2Av foci. (G) Wb of the experiment described in Figure 1D, shows the cytoplasmic fractions of indicated proteins. The Ponceau staining is shown as a loading control. (H) Quantification of the experiment described in Figure 1E shows the total number of γH2Av foci colocalizing with the indicated nucleoporins at different timepoints after IR. (I) Quantification shows Nup88, Nup98 or Sec13 foci colocalizing with γH2Av foci 30 min after IR, following indicated RNAi depletions. ****P<0.0001, n>193 foci for Nup88, n>75 foci for Sec13, and n>208 foci for GFP-Nup98FG. (J) Quantification of γH2Av foci colocalizing with H3K9me2 and Nup88, Nup98 or Sec13 at 10 min after IR. n>300 foci. Error bars are SEM in A,H and SD from three experiments in D-F. P values calculated with two-tailed Mann–Whitney test. Images are max intensity projections of a few Z-stacks. Scale bar= 1µm.

**Supplementary Figure 2. Relatated to figure 2. Constitutive anchoring of Nup88, Nup98, and Sec13 to the nuclear periphery results in defective relocalization.** (A) Wb analysis shows depletion efficiencies for Nup88 in Nup88-MycFLAG-Ndc1-expressing cells after Nup88 UTR RNAi, and for Sec13 in Sec13-MycFLAG-Ndc1-expressing cells after Sec13 UTR RNAi. (*) A non-specific band is used as loading control. (B-D) Quantifications of the experiments described in Figures 2a-c shows the total number of γH2Av foci at indicated timepoints after IR. (E-G) Schematic representations (top) show the experimental setup. Cells expressing wild type Nup88-MycFLAG (E) Nup98-MycFLAG (F) or Sec13-MycFLAG (G), were treated with RNAi directed to the corresponding endogenous protein (UTR RNAi). Quantifications of cells fixed 60 min after IR show the number of γH2Av foci in DAPI-bright and the total number of foci. n>173 cells for each RNAi condition. Error bars are SEM in B-D, and SD from three independent experiments in E-G.

**Supplementary Figure 3. Relatated to figure 1 and 2. Relocalization defects following nucleoporin RNAi or Ndc1 fusions is not a consequence of transport defects.** (A) Scheme of the light-induced nuclear transport assay LEXY (adapted from^42^). (B) Schematic representation of the experiment with the LEXY probe to measure nuclear pore export/import. Upon blue-light exposure, the NLS-mCherry-LEXY probe translocates from the nucleus to the cytoplasm, allowing the measure of export by live imaging. After blue-light exposure the probe returns to the nucleus, and import is measured by live imaging during a 30-min time interval as indicated. (C) Images and quantification of the nucleoplasmic/cytoplasmic ratio of fluorescence intensity over time, for cells treated with the drugs Kpt276 or Importazole (Ipz) (or DMSO control), as indicated. ****P<0.0001. n=75 cells/time point/condition. (D-F) Quantification of the nucleoplasmic/cytoplasmic ratio of fluorescence intensity over time for cells cells treated with the dsRNA targeting Nup88 (D, left), Nup98 (E, left), or Sec13 (F, left), or expressing Nup88-MycFLAG-Ndc1 (D, right), MycFLAG-Ndc1-Nup98 (E, right), Sec13-MycFLAG-Ndc1 (F, right), and also expressing the LEXY probe. *P<0.05, **P<0.01, ****P<0.0001. n=75 cells per each time point. (G,H) Quantifications show the number of γH2Av foci in DAPI-bright and total foci at indicated timepoints after IR, following indicated drug treatments. n>100 cells for each time point. Error bars are SEM in G,H, and SD from three experiments in C-F. P values calculated with unpaired T-test.

**Supplementary Figure 4. Relatated to figure 2. Nucleoplasmic Nup88, Nup98 and Sec13 are required for relocalization.** (A) Schematic representation of indicated WT and mutant proteins used in Figure 2. Numbers indicate aminoacid positions. (B) IF (left) shows Nup88 at the nuclear periphery in cells expressing FHA-Nup88FL following Nup88 UTR RNAi, and Nup88 released from the nuclear periphery in cells expressing FHA-Nup88ΔC following Nup88 UTR RNAi. Quantification (right) of the experiment described in Figure 2D shows the total number of γH2Av foci. (C) IF (left) shows Nup98 at the nuclear periphery in cells expressing FHA-Nup98FL following Nup98 UTR RNAi, and Nup98 released from the nuclear periphery in cells expressing FHA-Nup98FG following Nup98 UTR RNAi. Quantification (right) of the experiment described in Figure 2E shows the total number of γH2Av foci. (D) Schematic representation shows that Sec13 anchoring to the nuclear periphery is dependent on the N-terminal domain of Nup96, which complements the β-propeller of Sec13. Expression of a Nup96ΔN-MycFLAG mutant, while RNAi depleting endogenous Nup96, results in Sec13 release from the nuclear periphery. (E) Wb analysis shows Nup96 RNAi efficiency in cells expressing FHA-Nup98FL and Nup96ΔN-MycFLAG-expressing cells, following Nup96 RNAi depletion. dsRNAs for Nup96 target the region deleted in Nup96ΔN, which is present in endogenous Nup96. (F) IF (left) shows Sec13 at the nuclear periphery in cells expressing Nup96ΔN-MycFLAG following Ctrl RNAi, and Sec13 released from the nuclear periphery in cells expressing Nup96ΔN-MycFLAG following RNAi depletion of endogenous Nup96. Nup107 is not affected. Quantification (right) of the experiment described in Figure 2F shows the total number of γH2Av foci. (G) IF shows foci of Nup88 (left, staining with anti-HA antibodies) and Sec13 (right) colocalizing with γH2Av foci, in cells expressing Nup88ΔC or Nup96ΔN, respectively. Dashed boxes and zoomed details highlight colocalizations. Scale bars=1µm. Error bars are SEM in B,C,F.

**Supplementary Figure 5. Relatated to figure 3. Nup88, Nup98 and Sec13 act downstream from Smc5/6 to promote relocalization and to prevent abnormal Rad51 recruitment.** (A) qPCR analyses show Dhx9/Mle and MBD-R2 RNAi depletion efficiencies. (B) Quantifications of cells fixed 60 min after IR show the number of γH2Av foci in DAPI-bright and total foci following indicated RNAi depletions. n>363 cells. ****P<0.0001. P values calculated with two-tailed Mann–Whitney test. (C) Quantification of the experiment described in Figure 3b shows the total number of γH2Av foci. (D) Images (left) and quantifications (right) of indicated proteins in cells expressing GFP-Mu2, GFP-ATRIP, or GFP-Nse2 plus mCherry(mCh)-HP1a (heterochromatin mark^10^), as indicated, shows Nse2 protein enrichments in heterochromatin before IR and Mu2, Atrip, or Nse2 focus formation 10 min after IR, following indicated RNAi depletions. n>655 foci for Mu2, n>297 foci for ATRIP and n>155 foci for Nse2. Box plots: blue line=mean; black line=median. Whiskers are drawn down to the 10th percentile and up to the 90th from three experiments. (E) IF (top, example shown for Unc45) and quantification (bottom) of γH2Av foci colocalizing with FHA-Myo1A, FHA-Myo1B, FHA-MyoV, FHA-Arp3 or Unc45 10 min after IR, following indicated RNAi depletions. n>140 foci for each RNAi condition. (F) Quantification of dynamic nuclear actin filaments in response to IR in cells expressing the F-actin marker GFP-F-actCB-NLS, following indicated RNAi depletions, n>19 cells for each RNAi condition. (G) Quantification of foci after IF in undamaged cells (-IR) shows the number of Rad51 foci in DAPI-bright and total foci following indicated RNAi depletions. n>377 cells for each condition. (H) Quantification of the experiment described in Figure 4E shows the total number of Rad51 foci. Error bars are SEM in E, and SD in B,C,G,H from three independent experiments.

**Supplementary Figure 6. Relatated to figure 4. Nup88, Nup98, and Sec13 nucleoporins are required for mobilizing heterochromatic DSBs.** (A) LDM analysis of foci from the experiment described in Figure 4B. Duration and distribution of LDMs (blue lines) along each original track relative to different nuclear positions for each heterochromatic repair focus. (B) MSD analysis of time points corresponding to LDMs confirms that each LDM identifies a time interval characterized by directed motion, with the typical upward curvature of the graph^9^.

**Supplementary Figure 7. Relatated to figure 5. Nup98FG and Nup98YG phase separate *in vivo*, whereas Nup98AG does not.** (A) Analysis of Nup98 using the Predictor of Natural Disordered Regions (PONDR)^78^, shows the portions of the protein with higher predicted disorder. (B) Schematic representation of the constructs for the optoIDR assay (adapted from^55^). (C) Top: Schematic representation of the the optoIDR assay showing clustering of Cry2-mCherry fusion proteins into optoDroplets upon blue light exposure. Bottom: representative images taken at indicated time points from the end of the blue light exposure for cells expressing indicated constructs. Arrows and zoomed details highlight examples of droplet fusion (yellow) and splitting (magenta) events. See also Supplementary movie S1. (D) Quantification of the experiment in (C). n>13 cells for each condition. ****P<0.0001, **P<0.01 with two-tailed Mann-Whitney test from three or more experiments. Images (E) and quantification (F) show the reversibility and repeatable inducibility of Nup98FG optoDroplets by cycles of blue light activation for 4min followed by 30min recovery in the dark. n>42 cells/condition. Scale bars=1µm. In D and F, box plot: blue line=mean; black line=median. Whiskers are drawn down to the 10th percentile and up to the 90th from three experiments.

**Supplementary Figure 8. Relatated to figure 5. Nup98 mediates relocalization of heterochromatic DSBs *via* phase separation.** (A) Quantification of the experiment described in Figure 5B shows the total number of γH2Av foci. (B) Quantification shows the number of Rad51 foci in DAPI-bright and total foci in cells expressing FHA-Nup98-FG, -AG or -YG, following Nup98 UTR RNAi depletion in undamaged conditions (-IR), n>471 cells for each condition. (C) Quantification of the experiment described in Figure 5D shows the total number of Rad51 foci. (D) Quantification of cells fixed 60 min after IR show the number of γH2Av foci in DAPI-bright and total foci following indicated 1.6-Hexanediol treatments n>150 cells for each condition. (E) As in D, except Rad51 foci were quantified. n>462 cells for each condition. (F) Representative images from movies show Nup98AG condensates in cells expressing RGG-mCh-RGG-eDHFR and Halo-Nup98AG-GFP, after TNH addition to the media. The white arrowhead points to an example of signal overlap. (G) Quantification of the experiment described in Figure 5F shows the total number of γH2Av foci. (H) Quantification of the experiment described in Figure 5G shows the total number of Rad51 foci. Error bars are SEM in A and SD in B-E, G, H, from three or more experiments.

**Supplementary Figure 9. Relatated to figure 6. Nup88, Nup98 and Sec13 nucleoporins are required for heterochromatin repair and stability.** (A) IR sensitivity assay shows Kc cell survival at different IR doses and after indicated RNAi depletions. ****P<0.0001 for all comparisons *vs* Ctrl. n>280 cells/RNAi/dose. Rad51 RNAi is used as positive control. (B) Quantification of the experiment described in Figure 6c shows the number of persistent γH2Av foci not colocalizing with H3K9me2 signal (-H3K9me2). (C) Quantification of the experiment described in Figure 6d shows the number of micronuclei in undamaged conditions (-IR). +/− H3K9me2: micronuclei with/without H3K9me2. (D) Wb analyses show Nup98 depletion efficiency in Nup98 RNAi flies relative to ActinGal4/+ control (Ctrl), or the loss of Sec13 in *sec13^01031^* and *sec13^GS11919^* mutant flies relative to w^1118^ Ctrl, as indicated. Lamin or Tubulin are loading controls. qPCR analysis shows Nup88 depletion efficiency in Nup88 RNAi flies relative to ActinGal4/+ control (Ctrl). (E) IF shows normal Nup107 and nuclear pore protein staining (MAb414) in salivary glands and imaginal discs of Nup98 RNAi flies expressing Nup96-Myc. (F) Images of chromosome preparations from larval brains stained by FISH for AACAC, 359bp, and AATAT satellites and quantifications show chromosome fusions and abnormalities in Sec13 mutants, Nup88 and Nup98 RNAi *vs* control flies. In Nup98 RNAi, the arrowhead points to a translocation between chromosome X and Y, the asterisk highlights a chromosome fragment. In *Sec13^01031^,* the arrowhead highlights translocation between X and 4th, the dashed oval highlights an arm loss and the arrow a chromosome fusion between the two centromeric regions of chromosome 2. In *Sec13^GS11919^* arrowheads highlight fusions between chromosome 4 and the centromeric region of chromosome 3, and the arrow points to a filament connecting the centromeres of the two chromosome 2. *P<0.05, **P<0.01, ***P<0.001, ****P<0.0001, calculated with two-tailed Mann–Whitney test. n>58 karyotypes/genotype from at least two independent crosses and three brains. Scale bar =1µm. Error bars are SEM in F and SD from three experiments in A-C. Images are max intensity projections of a few Z-stacks.

**Movie S1. Nup98FG optoDroplet formation.** Cry2-mCherry, Cry2-mCherry-Nup98FG, Cry2-mCherry-Nup98AG, or Cry2-mCherry-Nup98AG was transiently expressed in Kc cells for 48 hours and clustering was induced by blue light exposure. Cells were imaged before, at 15 seconds followed by 45-s time intervals for 6 minutes. Representative cells corresponding to Figure S7C are shown. Scale bar 1µm.

## STAR METHODS

### Key Resources Table

Provided as a separate file

### Experimental model

#### Cell cultures

Kc167 (Kc) cells were used for all experiments and were maintained as logarithmically growing cultures in Schneider’s medium (Sigma) or Sf-900 II (ThermoFisher) + 10% FBS (GemCell US Origin, Gemini) + antibiotic-antimycotic (ThermoFisher). Kc cells were authenticated by the *Drosophila* Genomic Resource Center (DGRC) and no mycoplasma contamination was detected.

#### Flies

All crosses were carried out at 25°C, and flies were maintained as previously described^79^. Stocks were obtained from: the Vienna *Drosophila* Resource Center (VDRC) (Nup88 RNA: stock #v22446); the Bloomington *Drosophila* Stock Center (BDSC) (Sec13^01031^ mutant: #10327); the Kyoto *Drosophila* Genetic Resource Center (DGRC) (Sec13^GS11919^ mutant: #203684); and the Capelson Lab (Nup98RNAi;Nup96^50^). *w^1118^* was a kind gift from J. Tower. Act5c-GAL4 was from Bloomington (#25374). See also Key Resources Table.

### Method Details

#### IR Treatments

Cultures were exposed to IR using a 160 kV X-ray source (X-RAD iR-160, Precision X-Ray). We use a range of Gy at which the damage response (estimated based on the number of γH2Av foci) increases linearly with dose, and corresponds to nearly sublethal doses for controls (more than 90% survival at 2.5-10 Gy). A dose of 5 Gy was used for most experiments, unless otherwise indicated. The estimated number of DSBs induced by 5 Gy in *Drosophila* cells is approximately 7.5 DSBs in G1 and 14 DSBs in G2, which is based on published estimates of DSB numbers in mammalian cells, and the estimated genome size for human vs *Drosophila* cells. In kinetic analyses of fixed cells, time 0 (Unt) corresponds to cells fixed without exposure to IR. In time-lapse experiments, time 0 (Unt) corresponds to cells imaged 5 min before IR treatment.

#### Generation of cell lines expressing tagged proteins

Most experiments were performed using stable cell lines, obtained by cotransfecting the plasmid(s) of interest with pCoHygro (Invitrogen) or pCoPuro (Addgene plasmid #17533) and selecting in the presence of 100 µg/ml Hygromycin B (Enzo Life Sciences) or 2 mg/ml Puromycin (Sigma-Aldrich). Transfections were performed with Cellfectin (Life Technologies), according to manufacturers’ procedures. For transient transfections (Figure S7C and Supplementary movie S1), cells were split 2 days before transfection in 1ml-wells, and 3×10^6^ cells were transfected with 2.5 ug of plasmid DNA, using Cellfectin (Invitrogen) following manufacturer’s instructions. Imaging was done 3 days after transfection.

#### Plasmids

See additional information for plasmids in Key Resources Table. pCopia-mCherry(mCh)-HP1a, pCopia-GFP-Mu2, pCopia-GFP-ATRIP, pCopia-GFP-Nse2, pCopiaF-actCB–GFP–NLS plasmids were previously described^10,13^. All other FHA-tagged (C-terminus) or MycFLAG-tagged (N-terminus) full-length (FL) proteins were generated by insertion of PCR-amplified sequences using the following cDNA from DGRC as templates: Nup88 (LD33348), Sec13 (LD03471), and Nup98-96 (RE74545). Nup88, and Nup98 PCR products were cloned into AscI/PacI-digested pCopia-3xFLAG-StrepII-3xHA (FHA) vector (described in^10^). Nup96, Nup88 and Sec13 were cloned into an KpnI/AccIII-digested pCopia-Myc-StrepII-3xFLAG vector (kind gift from G. Karpen). Nup98FL and Nup96FL were generated by PCR amplification of the corresponding sequences of the Nup98-96 fusion template.

Ndc1 fusions were created by PCR-amplification of the *Drosophila* Ndc1 cDNA as template (DGRC, LD42327). Specifically, Nup88-MycFLAG-Ndc1 and Sec13-MycFLAG-Ndc1 plasmids were generated by subcloning the coding region of Ndc1 into KpnI/AccIII-digested pCopia-Nup88-MycFLAG and pCopia-Sec13MycFLAG plasmids, respectively. pCopia-MycFLAG-Ndc1-Nup98 plasmid was generated by subcloning the MycFLAG-Ndc1 sequence from pCopia-Sec13MycFLAG-Ndc1 plasmid into Eco47III/AscI-digested pCopia-FHA-Nup98 plasmid. pCopia-Nup96ΔN-MycFLAG was generated by subcloning the PCR-amplified Nup96ΔN sequence into AscI/PacI-digested pCopia-Nup96-MycFLAG. pCopia-FHA-Nup88ΔC was generated by subcloning the PCR-amplified Nup88ΔC sequence into AscI/PacI-digested pCopia-FHA-Nup88, and an SV40 NLS was inserted in C-terminus to compensate for the loss of a predicted NLS sequence in the Nup88ΔC mutant^47^.

pCopia-FHA-Nup98FG was generated by subcloning the Nup98FG sequence from DamNup98FG plasmid (kind gift from M. Fornerod^30^), into AscI/PacI-digested pCopia-3XFLAG-StrepII-HA vector. pCopia-FHA-Nup98AG, pCopia-FHA-Nup98YG, were generated by PCR amplification of Nup98AG and Nup98YG (obtained as geneblocks from GENEWIZ) and cloning into AscI/PacI-digested pCopia-3xFLAG-StrepII-3xHA vector.

To generate pAWG-GFP-Nup98AG and pAWG-GFP-Nup98YG plasmids, Nup98AG and Nup98YG were subcloned into SacII/AscI-digested pAWG-EGFP-Nup98FG vector (kind gift of M. Hetzer).

To adapt the LEXY system to *Drosophila* cells, the mCherry-LEXY (AsLOV2-NES21) fragment was PCR amplified from pDN122 (pCMV-NLS-mCherry-LEXY from Barbara Di Ventura & Roland Eils^42^, Addgene plasmid #72655), and cloned into a pCopia backbone (pCopia-eGFP-Arp3^13^) after NheI/PacI digestion. The *Drosophila-*codon-optimized “weak” NLS sequence KRKYF^80^ was also included in N-terminus through the cloning oligos, in place of the original cMyc^P1A^ NLS^42^. In our initial tests, the cMyc^P1A^ NLS did not drive the nuclear localization of the LEXY construct to the nucleus with high efficiency, while the “class 4” (KRKYF) NLS did.

For Cry2-containing plasmids, the N-terminal PHR domain of the *A. thaliana* photoreceptor cryptochrome 2 (Cry2PHR) was PCR amplified with the N-terminal mCherry from pCRY2PHR-mCherryN1 (from Chandra Tucker, Addgene plasmid # 26866^81^), and cloned into a pCopia-Flag backbone to generate pCopia-Flag-Cry2PHR-mCherry. pCopia-Flag-Cry2PHR-mCherry-Nup98FG/AG/YG plasmids were generated by subcloning the Nup98FG/AG/YG sequence from pCopia-FHA-Nup98FG/AG/YG plasmids into AscI/PacII-digested pCopia-Flag-Cry2PHR-mCherryN1.

pCopiaRGG-mCherry-RGG-eDHFR was generated by PCR amplification of RGG-mCherry-RGG-eDHFR from pHZ088-RGG-mCherry-RGG-eDHFR (kind gift from H. Zhang and M. Lampson^59^) and cloning into a NheI/PacI-digested pCopia-3xFlag-StrepII-3xHA vector. pCopia3xHalo-GFP was generated by PCR amplification of 3xHalo-GFP from pHZ031-3xHalo-GFP (kind gift from H. Zhang and M. Lampson^59^) and cloning into NheI/PacI-digested pCopia-3xFLAG-StrepII-HA. pCopia3xHalo-GFP-Nup98AG was created by PCR amplification of GFP-Nup98AG from pCopia-FHA-Nup98AG, and cloning into AscI/PacI-digested pCopia-3xFLAG-StrepII-HA.

#### dsRNA synthesis and sequences

dsRNAs were prepared with the MEGAscript T7 Kit (Thermo Fisher Scientific Cat# AM1334). Amplicons for Bw, Smc5, Smc6, Rad51, Nse2/Cerv, and Nse2/Qjt dPIAS dsRNAs were previously described^6,10,12^. Amplicons used for all other dsRNAs were: Nup88 (DRSC16123), Sec13 (DRSC14114), Elys (DRSC31834/DRSC19571), Nup214 (DRSC04421/DRSC39709), Nup62 (DRSC06970/DRSC34583), Nup358 (DRSC34580/DRSC29772), MDB-R2 (DRSC37591/DRSC32494), Dhx9 (DRSC04921/DRSC37605). Sequences can be found on the DRSC website (http://flyrnai.org). The amplicon for Nup98 was previously described (see Table S1 and^29^). dsRNAs for Nup88UTR, Sec13UTR, Nup98UTR, and Nup96 RNAi depletions were prepared with the oligos listed in Table S1. When more than one amplicon is indicated, we combined equal amounts of each dsRNA for better efficiency of protein depletion.

#### RNAi depletions in cultured cells

dsRNAs were transfected with DOTAP Liposomal Transfection Reagent (Roche) following manufacturer’s instructions. Incubation times and dsRNA amounts were optimized to maximize depletion efficiency while avoiding toxicity and cell cycle effects. For most RNAi depletions, cells were treated with dsRNAs for 5 days, except: Nup88 or Nup88 UTR RNAi was done for 6 days; Sec13 RNAi was done for 6 days; Nup98 RNAi was done for 3 days; RNAi depletion of endogenous Nup96 was done for 4 days. The control (Ctrl) used for all RNAi experiments is RNAi depletion of the *brown (bw)* gene, which regulates the body color of adults flies and is not involved in DNA repair pathways^10,12,13^. RNAi depletion efficiencies for *bw*, Smc5/6, Nse2/Cerv, Nse2/Qjt, dPIAS, Rad51, have been previously validated^10,12,13^. RNAi depletion of endogenous Nup96 in Nup96ΔN-MycFLAG-expressing cells is always done in the presence on FHA-Nup98FL, because the amplicon targeting Nup96 also affects the level of Nup98 it is coexpressed with^82^. For the same reason, all RNAi depletions of Nup98 are done in cells expressing Nup96-MycFlag. We note that all kinetics resulting from RNAi depletion must be compared to cells treated with control dsRNAs, since the γH2Av peak shifts from 10 min after IR in non-RNAi experiments to 30 min after IR in RNAi controls^10^.

#### Antibodies

Antibodies used were: anti-Nup88 (1:1000 for IF, 1:10000 for Wb, kind gift from M. Hetzer); anti-Sec13 (anti-Sec13 Guinea Pig used at 1:1000 for IF; anti-Sec13 Rabbit, used at 1:200 for Wb, M. Capelson); anti-Nup98 (1:1000 for IF, Capelson lab); anti-Nup62 (1:1000 for IF, Ohkura lab); anti-Nup107 (1:2000 for IF, kind gift from V. Doye), anti-FLAG (1:500 for IF, 1:1000 for Wb, Sigma); anti-HA (1:1000 for IF, Developmental Studies Hybridoma Bank); anti-GFP (1:500 for IF, Aves Lab); anti-H3K9me2 (1:500 for IF, Wako Chemicals); anti-H3K9me3 (1:500 for IF, Wako Chemicals); anti-H4 (1:1000 for Wb, Sigma); anti-Myc (1:1000 for IF, Developmental Studies Hybridoma Bank); anti-γH2Av (1:500 for IF, Rockland); anti-Rad51 (1:500 for IF, gift from J. Kadonaga); anti-Unc45 (1:500 for IF, gift from S. Bernstein); anti-Lamin/Dm0 (1:1000 for IF, 1:2000 for Wb, Developmental Studies Hybridoma Bank); anti-β-actin (1: 2500 for Wb, Abcam); anti-α-tubulin (1:1000 for Wb, Sigma); anti-nuclear pore complex proteins (MAb414, 1:1000 for IF, Biolegend). Additional information on antibodies is provided in Key Resources Table. Secondary antibodies for immunofluorescence were from Life Technologies and Jackson Immunoresearch. Those used for western blotting were from Pierce and Santa Cruz Biotech. Antibodies were previously validated^10,12,13^ or validated by comparing western blot or immunofluorescence signals in the presence of the protein of interest with signals after RNAi depletions, or immunofluorescence signals in the absence of primary antibodies.

#### qPCR

Total RNA was isolated from 3-5 × 10^6^ cells or 5-10 larvae by Trizol extraction and treated with DNaseI to remove any genomic DNA contaminant. RNA was used to generate single-stranded cDNA using random priming and Superscript Reverse Transcriptase III (Invitrogen) or SuperScript IV stand-alone reverse transcriptase (Thermo Fisher Cat# 18090050). Specific transcripts were quantified using intron-spanning primers with iQ SYBR Green Supermix (Bio-Rad) and using a MyiQ Single Color, Real-Time PCR Detection System (Bio-Rad), according to manufacturer’s instruction. Changes in transcript levels were normalized to Act5C mRNA for qPCR analysis of transcripts in cultured cells and to Lamin mRNA for qPCR analysis of transcripts in larvae. Each qPCR was repeated at least three times, and graphs show the average level of depletion relative to control RNAi. Oligonucleotides used for qPCR are listed in Table S1.

#### Western blotting

1-3×10^6^ cells were collected, washed once in PBS and lysed for 15–20 min on ice with lysis buffer A (50 mM Tris, pH 7.8, 1% NP-40, 150 mM NaCl) containing protease inhibitors (Complete, Roche), 2-mercaptoethanol, and 1 mM PMSF. Benzonase was added to each sample (0.5U/ul) for 30 min. The soluble lysate was recovered by centrifugation (10 min, 4°C) and resuspended in the loading buffer (Laemmli) to a final concentration of 1x. Samples were denatured for 5 min at 95°C before running them on a TGX 4–12% or 10% gel (Bio-Rad) and transferred onto nitrocellulose membrane for hybridization with specific antibodies.

#### Chromatin fractionation

Fractionation of Kc cells was performed using the Thermo Scientific Subcellular Protein Fractionation kit for cultured cells (cat# 78840), following manufacturer’s instructions. The quality of fractionation was validated using antibodies specific for the different fractions.

#### Immunofluorescence and quantification of Repair Foci in Fixed Samples

Immunofluorescence without triton extraction was used for most experiments as previously described^10,12^. Immunofluorescence staining for Figure 1E, 3A, 3C, 5C and Figure S1I, 4G, 5E was preceded by a triton extraction step as previously described^12,13^, which removes most of the non chromatin-bound and nuclear periphery-associated signal of nucleoporins. Imaging and image processing for fixed cells and tissues with the DeltaVision Deconvolution microscope and the SoftWorx Software has previously been described^10,12^. In IF experiments, classification of repair foci inside or outside the DAPI-bright region was done as previously described^10,12^. Classification of foci inside and outside the heterochromatin domain was done by analyzing the position of foci relative to the H3K9me2 staining in each of the Z-stacks. Foci associated with the heterochromatin domain were either inside the H3K9me2 domain, at the periphery of the domain, or at the tips of H3K9me2 protrusions^10,12^.

#### Cell Imaging and Processing in Time-Lapse Experiments

Time-lapse experiments and quantifications were performed as previously described^12,13,54,83^. For MSD analyses, cells were imaged with 40-sec time intervals for 60 min starting from 3-5 min after IR. 10 Z-stacks at 0.8 µm distance were imaged for 0.15 ms for GFP, and 0.015 ms for mCherry with 10% transmission light, using a DeltaVision microscope (GE Healthcare/Leica) and an Olympus PlanApo 60X oil objective N.A. 1.42. The Coolsnap HQ2 camera was set at 2×2 binning for maximizing the intensity of the light collected and minimizing light exposure. All movies were corrected to compensate for modest photobleaching effects using SoftWorks (Applied Precision/GE Healthcare). For each nucleus, foci that appeared mostly “static” were tracked with a semi-automated method in Imaris (Bitplane) and the “correct drift” function was applied to use these tracks for registering the nucleus. Foci were then tracked in 3D using an automated method and manually corrected to ensure optimal connections between timepoints. Focus positional data were extracted in Excel and analyzed in Matlab (MathWorks) and R using customized scripts^54^. MSDs were calculated as described in^54^. Positional data were also analyzed using a customized script in R Studio to detect LDMs^54^. LDMs shown in Figure 4B and Figure S6A correspond to the largest contiguous time interval containing MSDs of increasing slope for each focus. The ability of the script to correctly detect directed motions was confirmed by independent MSD analysis of the positional data within the time intervals of the LDMs, as shown in Figure S6B. For live imaging of nuclear actin filaments, a stable cell line expressing F-actCB–GFP–NLS was used^13^. The same field of cells was imaged before and every 5 min after IR (starting 15 min after IR) for 55 min. Ten *Z* stacks at 0.8 µm distance were imaged. 3D volume reconstructions and movie generation were done using Imaris (Bitplane).

#### Light-induced nuclear pore transport assay

Nuclear transport assay was adapted from^42^. Cells were imaged in the mCherry channel to establish the initial nuclear signal intensity, using 2×2 binning as described above with the CoolSnapHQ2 camera mounted on a DeltaVision microscope. Next, cells were exposed to blue light (excitation 488nm), through 1s-pulses every 30s for 15 min, which triggered the nuclear export of the LEXY construct. Cells were then imaged at 1, 5, 15, and 30 min during recovery. 10 Z-stacks at 0.8 µm distance were imaged for 0.08 ms with 32% transmission light for visualization of the mCherry signal, using a DeltaVision deconvolution microscope (GE Healthcare/Leica) and an Olympus PlanApo 60X oil objective N.A. 1.42. Light intensity from the nucleus and cytoplasm was quantified in the middle Z-stack for each cell using SoftWoRx or ImageJ and compared across timepoints. When indicated, nuclear transport was inhibited by treating cells with 100 µM Kpt276 (Sellek, Cat# S7251) or 20 µM Importazole (Ipz; Sellek, Cat #S8446) 40 min prior to imaging.

#### Cry2 optodroplet Assay

Cells used for this assay were kept wrapped in aluminum foil to limit light exposure before the experiment. On the day of the experiment, cells were imaged once, then exposed to blue light through 1s-pulses every 25s for 4 min. Cells where then imaged with 45-sec time intervals for 6 min starting from 15s after blue light exposure. 10 Z-stacks at 0.8 µm distance were imaged for 0.08 ms with 32% transmission light ms for visualization of the mCherry signal. The Coolsnap HQ2 camera was set at 3×3 binning for maximizing the intensity of the light collected and minimizing light exposure. All movies were corrected to compensate for modest photobleaching effects using SoftWorks (Applied Precision/GE Healthcare). Acquisitions were done using a DeltaVision deconvolution microscope (GE Healthcare/Leica) and an Olympus PlanApo 60X oil objective N.A. 1.42. For Figure S7E, optodroplets were induced in cycles through: blue light exposure, imaging of mCherry, incubation in the dark for 30 min, before the new cycle.

#### 1,6-Hexanediol treatments

2.5, 5 or 7.5% 1,6-Hexanediol (Sigma-Aldrich, Cat #240117) was added to the medium of *Drosophila* Kc cells starting from 10 min before 5 Gy IR exposure. Cells were fixed 1h after IR.

#### Dimerization Assay

To artificially fuse two arginine/glycine-rich (RGG) domains from the P granule component LAF-1 to Nup98AG, cells transfected with RGG-mCherry-RGG-eDHFR and 3xHalo-GFP-Nup98AG plasmids were treated with the dimerizer TNH: TMP-NVOC-Halo^58,59^. For live imaging, 1 µM TNH was added directly to and mixed with cells in a chambered coverslip, between imaging timepoints. For IF, 1 µM TNH was added to the cell media and incubated for 2h before fixing. Samples were covered to minimize exposure of the dimerizer to light before the experiments. Details of dimerizer synthesis and characterization with NMR spectra were previously described^58^.

#### IR-sensitivity assay

IR sensitivity assay for RNAi depletion experiments in Figure S9A was performed as previously described^13^. IR sensitivity assays for Ndc1-tagged Nup88 or Sec13 was performed as follows: dsRNAs for Nup88 or Sec13 UTRs were added on day 0 and 3, cells were irradiated on day 5, and quantified using trypan blue staining on day 8. IR sensitivity assays for Ndc1-Nup98 and Nup98FG were performed as follows: Rad51 dsRNAs were added on day 0, and Nup98UTR dsRNAs were added on day 2; cells were irradiated on day 5 and quantified on day 6. IR sensitivity assays for “nucleoplasmic-only” Sec13 were performed as follows: Sec13UTR dsRNAs were added on day 0 and 3, Rad51 dsRNAs were added on day 1 and 3, endogenous Nup96 dsRNAs were added on day 2. Cells were irradiated on day 6, and quantified on day 8. IR sensitivity assays for Nup88ΔC were performed as follows: dsRNAs for endogenous Nup88 were added on day 0 and 4, Rad51 dsRNAs were added on day 1 and 4. Cells were irradiated on day 6, and quantified on day 9. IR sensitivity assays for the Nup98AG mutant were performed as follows: Nup98 UTR dsRNAs were added on day 0 and 3. Cells were irradiated on day 5, and quantified on day 6.

#### Micronucleus assays

For detecting micronuclei, Nup88 and Sec13 dsRNAs were added on day 0, and 5, and Nup98 dsRNAs were added on day 3 and 5. Cells were irradiated with 10 Gy on day 6, fixed on day 9 and processed for IF analysis. Micronuclei were detected as DAPI-positive signals outside the nuclear periphery. Most micronuclei are “erupted”, *i.e.,* they do not show an intact nuclear membrane.

#### Fly crosses and Karyotyping by FISH

To obtain 3rd instar larvae for karyotyping of neuroblast metaphase spread, RNAi lines (See Key Resources Table.) were crossed to the Act5c-GAL4 line and non-GFP larvae were picked for karyotyping as described^84^. Chromosome preparation and FISH protocols were previously described^12,13,84^. Labeled AACAC, AATAT, and 359 bp probes were purchased from Integrated DNA Technologies. FISH probe sequences are listed in Table S1.

#### Statistical analysis

All statistical analyses were performed using Prism6 (Graphpad Software) or Excel.

## Lead Contact and Materials availability

Further information and requests for resources and reagents should be directed to and will be fulfilled by the lead contact, Irene Chiolo.

## Acknowledgments

This work was supported by AAUW International Fellowship to C.M.; USC Research Enhancement Fellowships to T. Ryu; USC Rose Hills Foundation Fellowship to N.B., NIH Training grant to C.S.; NIH R01GM124143 to M.C.; R01GM117376 and NSF Career 1751197 to I.C. We thank all the members of the Chiolo Lab for helpful discussions; G. Karpen, M. Fornerod, M. Dasso, B. Di Ventura, R. Eils, C. Tucker, M. Hetzer, M. Lampson, and H. Zhang for plasmids; Alexei Arnaoutov from M. Dasso Lab for helpful advice; M. Hetzer, H. V. Doye, J. Kadonaga, H. Ohkura and S. Bernstein for antibodies; J. Tower for fly lines and media; BDSC (P40OD018537), VDRC, and DGRC (P40OD010949) for flies and plasmids. A. Christensen for input on the manuscript. L. Boninsegna for help with LDM analyses, Rc calculations, and input on the manuscript. A. Patel and L. Oei for generating constructs. G. Dialynas for help with larval tissue staining. *Drosophila* Rad51 antibodies are now maintained and distributed by the Chiolo Lab. Antibodies against Lamin, HA and Myc were obtained from DSHB, created by the NICHD of the NIH and maintained at the University of Iowa.

