## Supplementary figures and images for "“Off-pore” nucleoporins relocalize heterochromatic breaks through phase separation"

### Supplementary figures

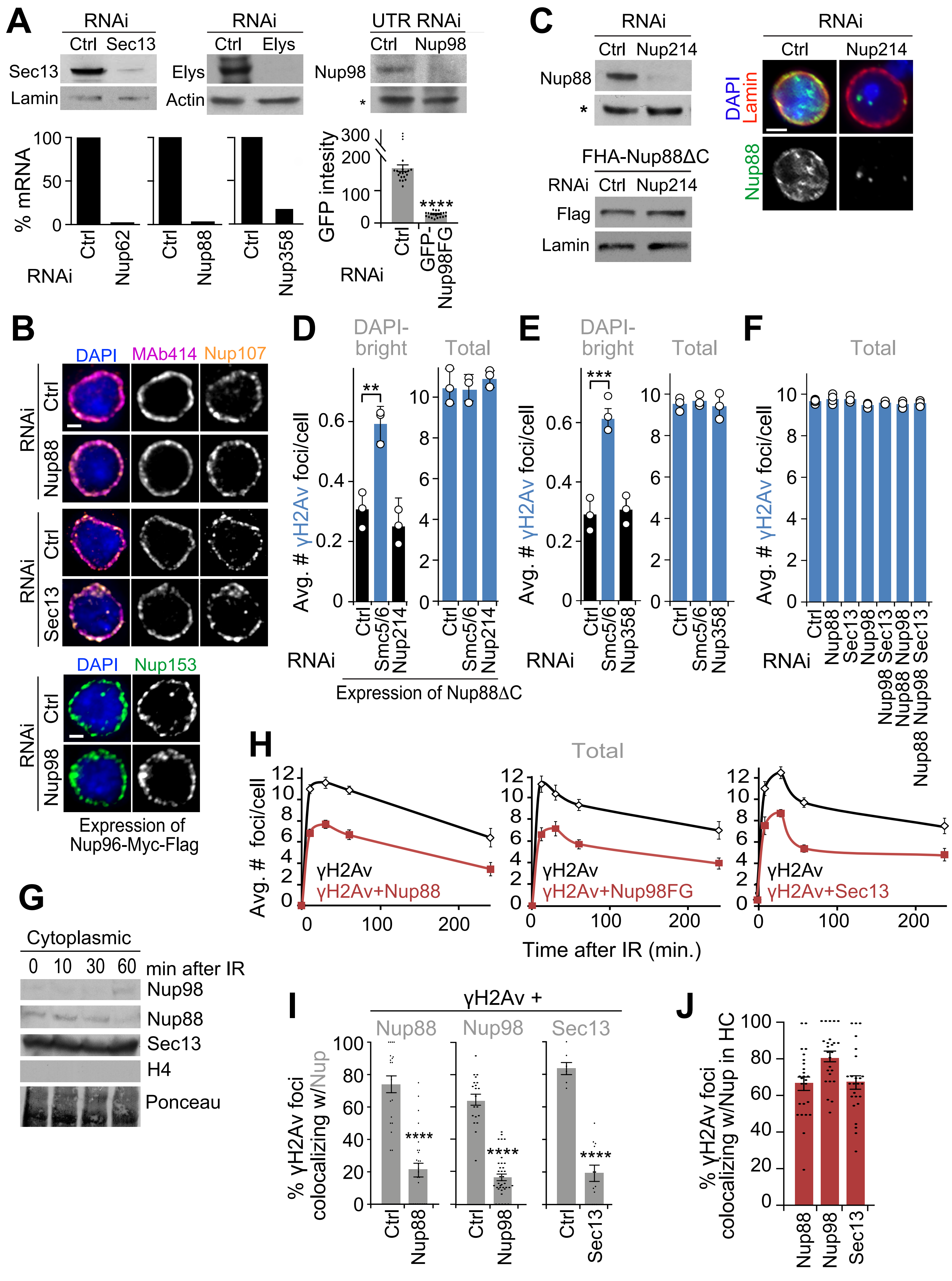

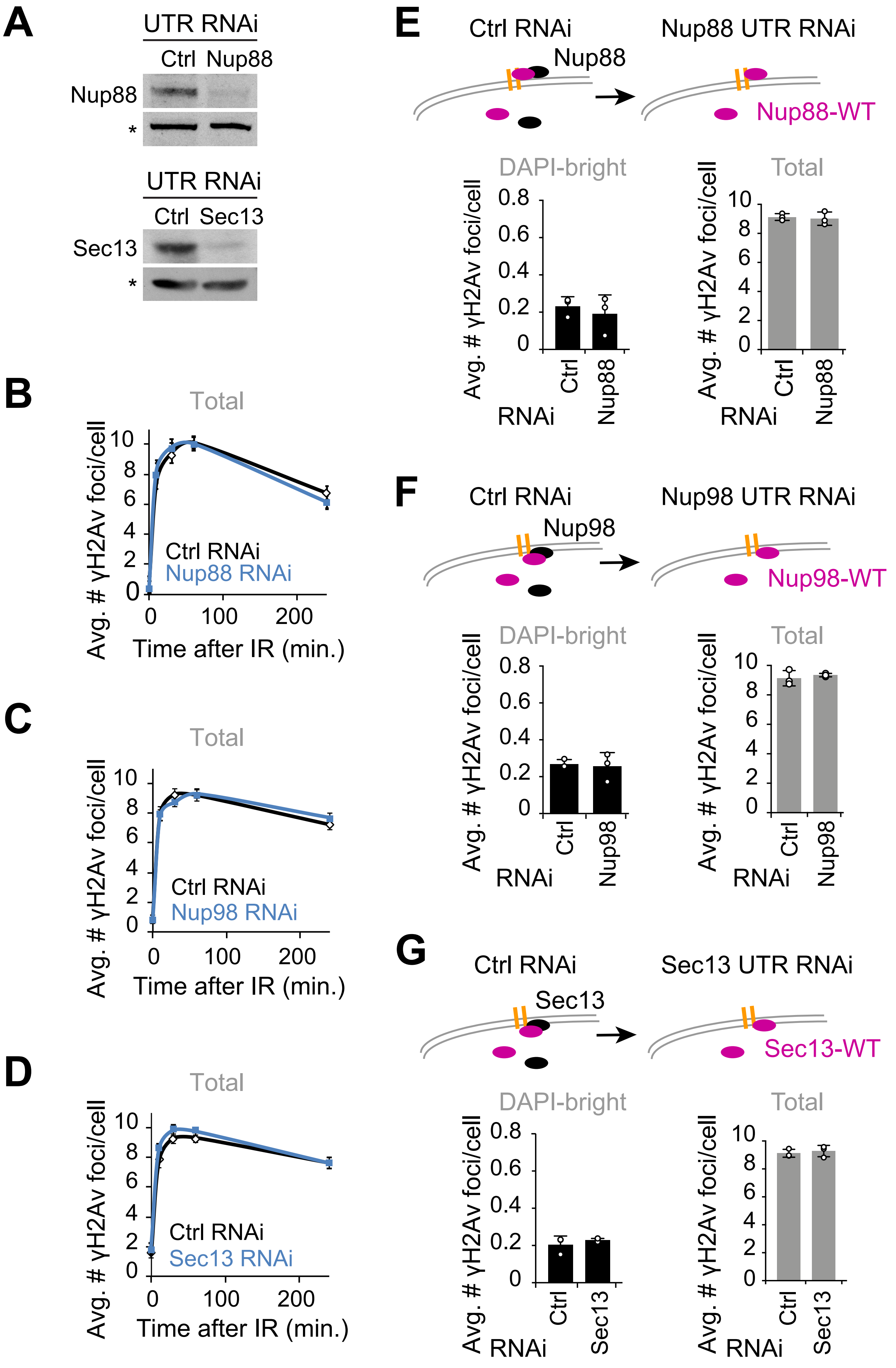

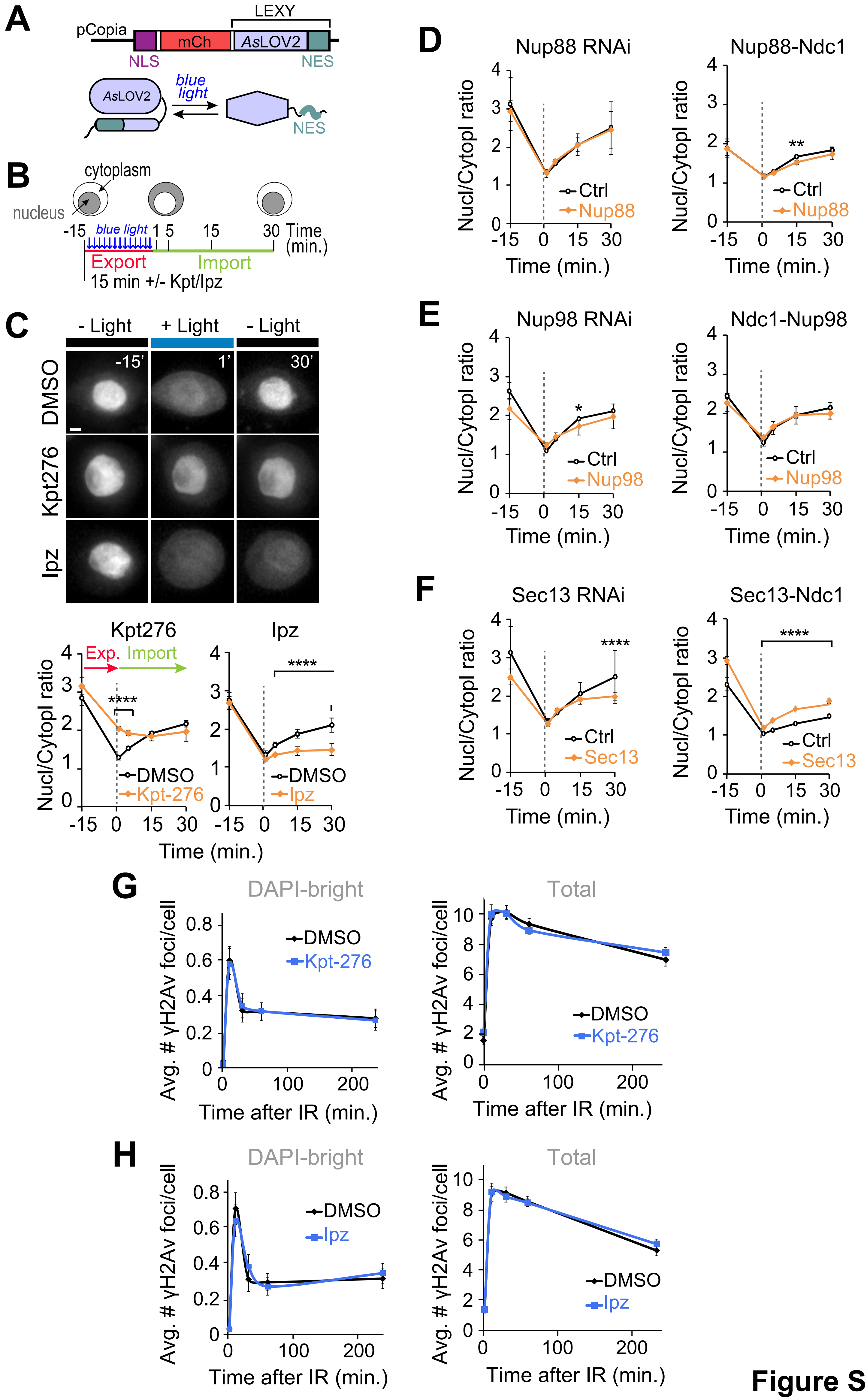

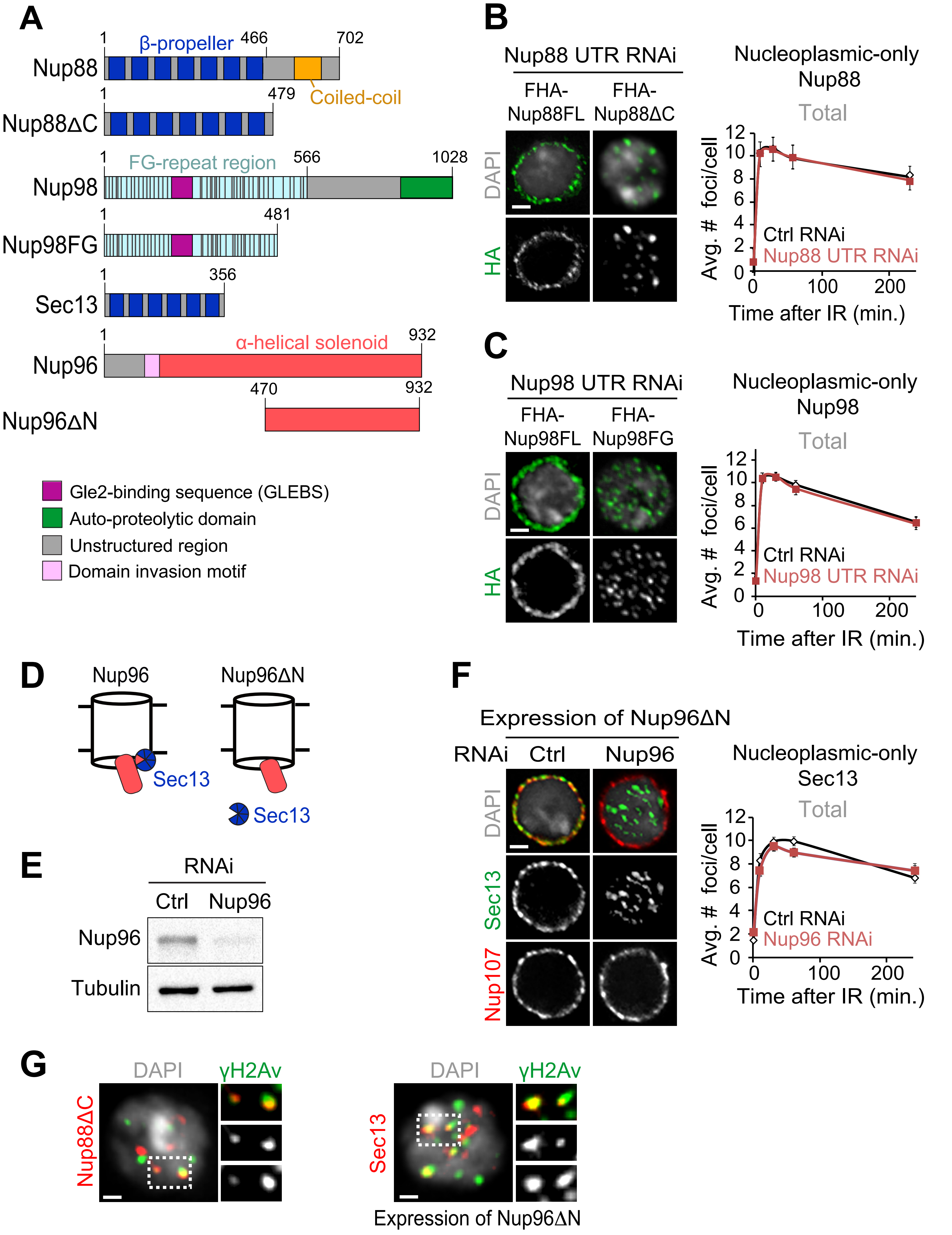

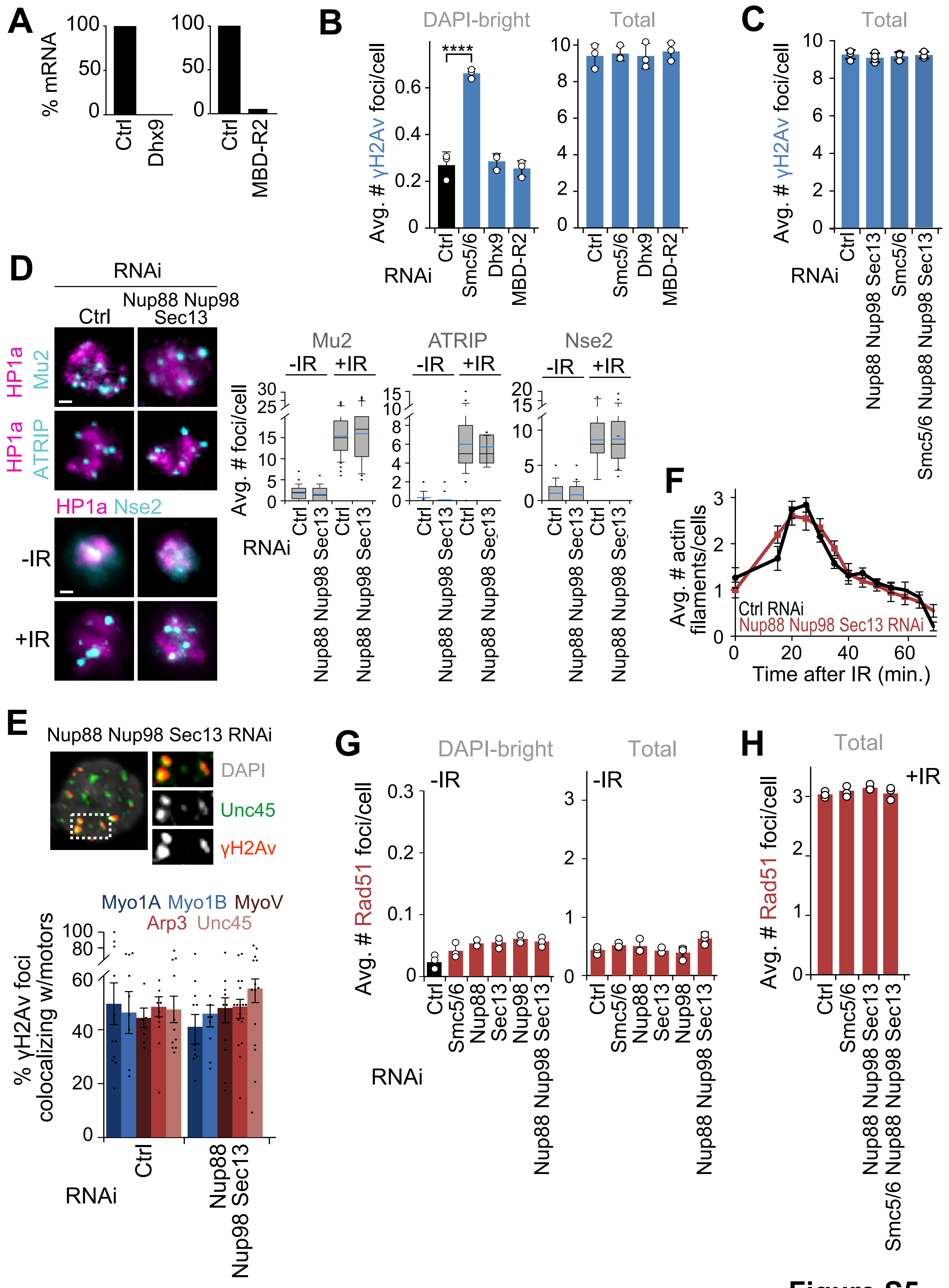

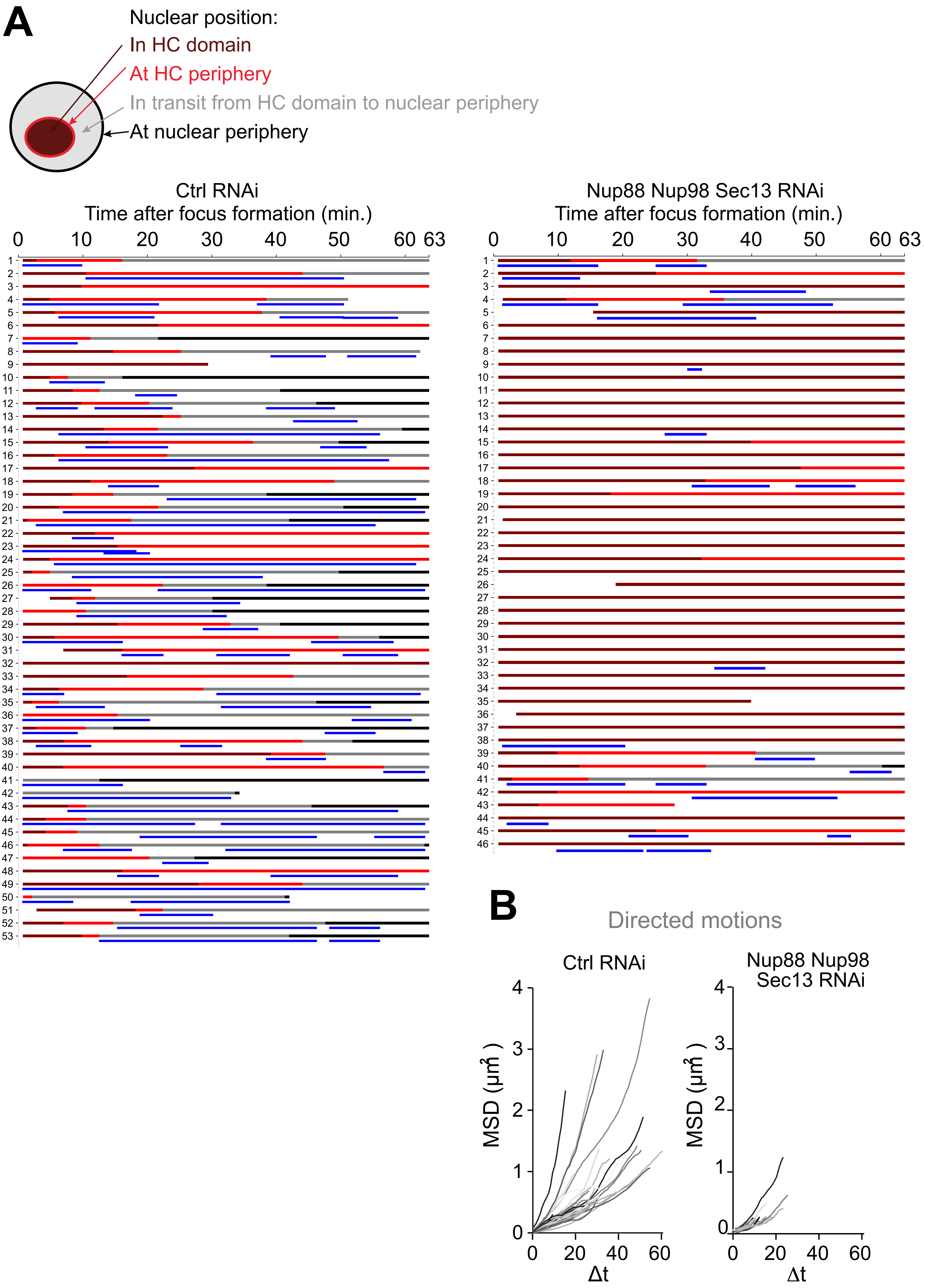

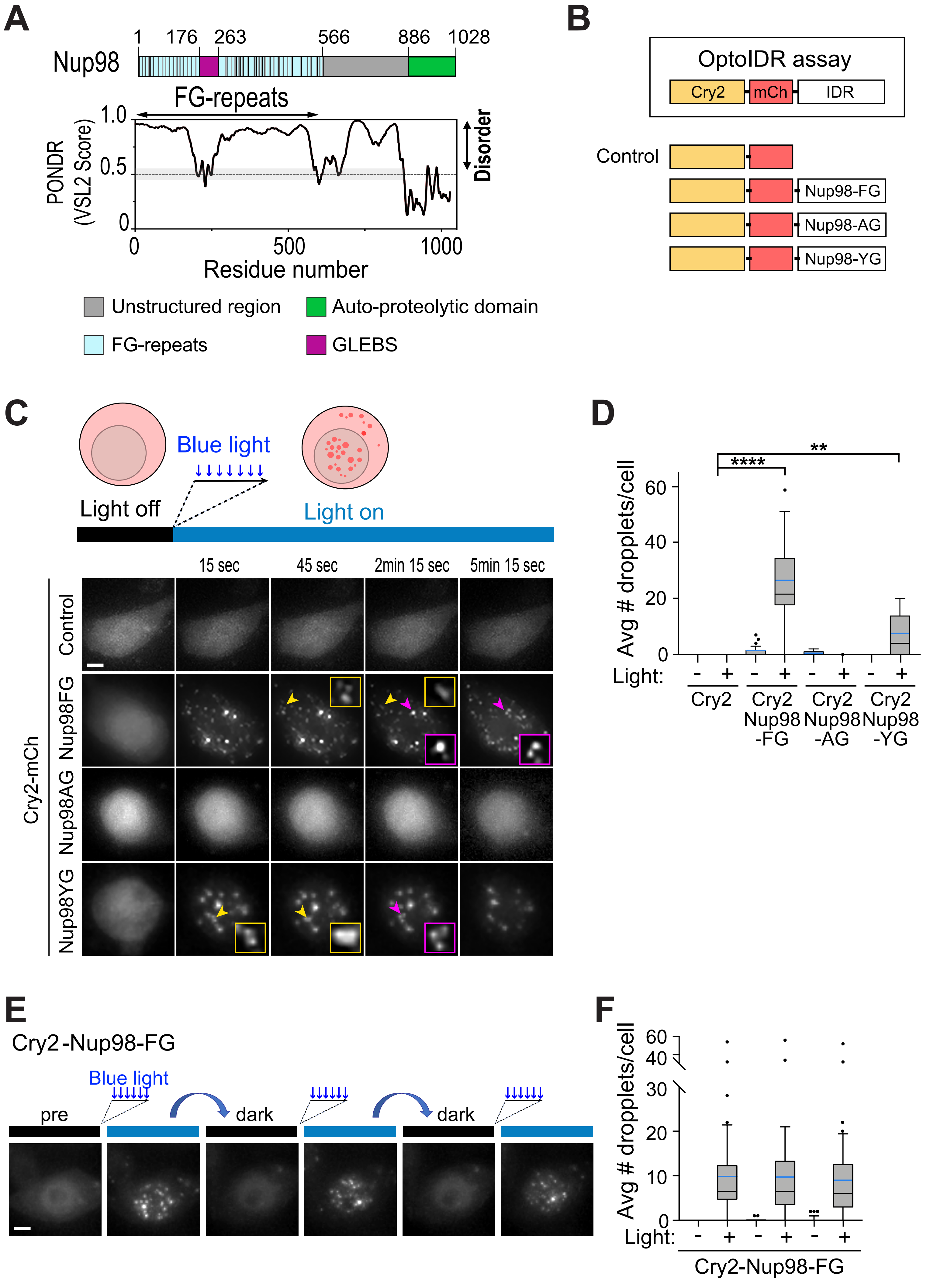

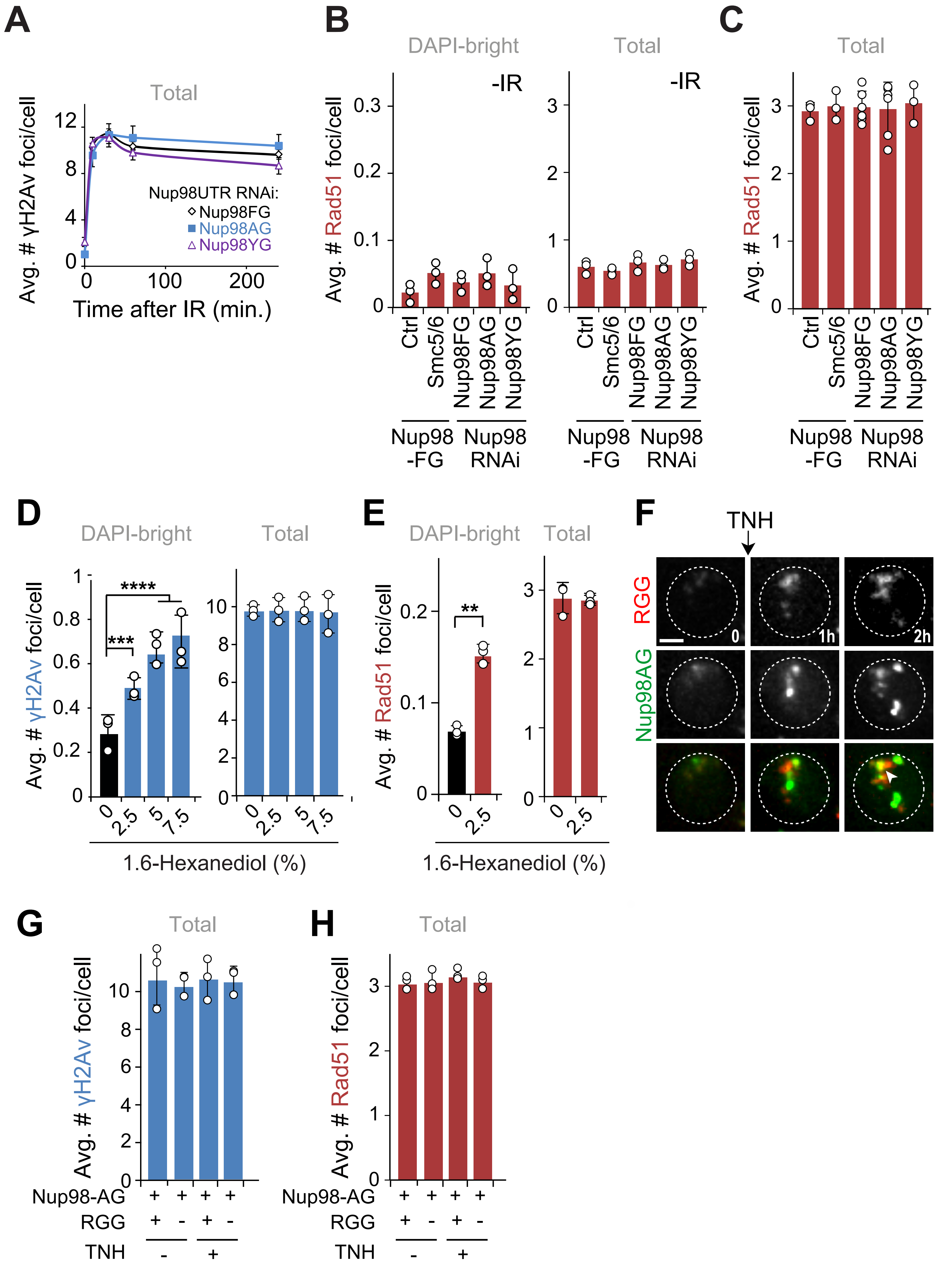

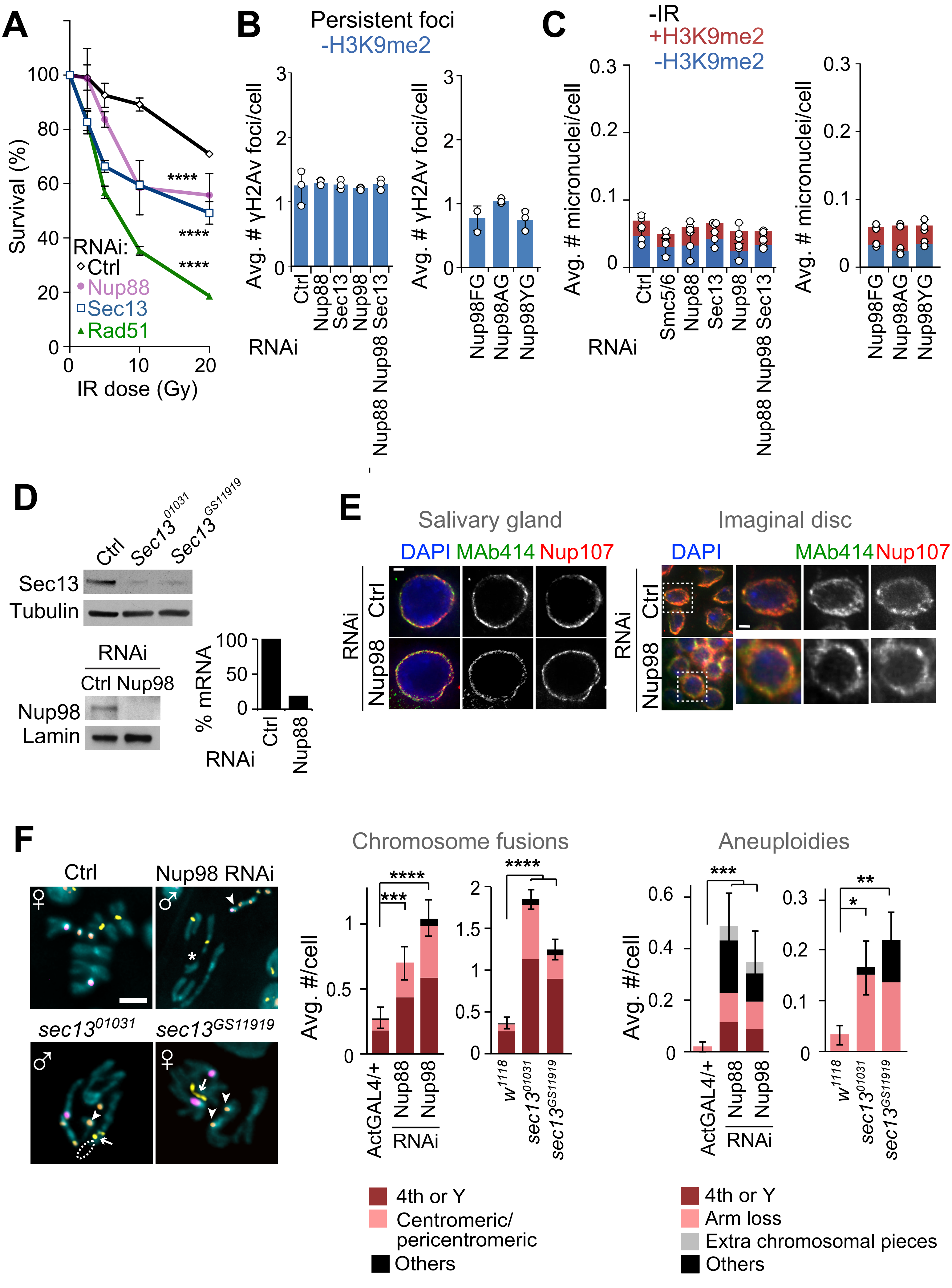
